# Pooled clone collections by multiplexed CRISPR-Cas12a-assisted gene tagging in yeast

**DOI:** 10.1101/476804

**Authors:** Benjamin C. Buchmuller, Konrad Herbst, Matthias Meurer, Daniel Kirrmaier, Ehud Sass, Emmanuel D. Levy, Michael Knop

## Abstract

Clone collections of modified strains (‘libraries’) are a major resource for systematic studies with the yeast Saccharomyces cerevisiae. Construction of such libraries is time-consuming, costly and confined to the genetic background of a specific yeast strain. To overcome these limitations, we present CRISPR-Cas12a (Cpf1)-assisted tag library engineering (CASTLING) for multiplexed strain construction. CASTLING uses microarray-synthesized oligonucleotide pools and in vitro recombineering to program the genomic insertion of long DNA constructs via homologous recombination. One simple transformation yields pooled libraries with >90% of correctly tagged clones. Up to several hundred genes can be tagged in a single step and, on a genomic scale, approximately half of all genes are tagged with only ∼10-fold oversampling. We report several parameters that affect tagging success and provide a quantitative targeted next-generation sequencing method to analyze such pooled collections. Thus, CASTLING unlocks new avenues for increased throughput in functional genomics and cell biology research.

## Introduction

The systematic screening of arrayed biological resources in high-throughput has proven highly informative and valuable to disentangle gene and protein function. For eukaryotic cells, a large body of such data has been obtained from yeast strain collections (‘libraries’) in which thousands of open reading frames (ORFs) are systematically altered in identical ways, e.g. by gene inactivation or over-expression to determine gene dosage phenotypes and genetic interactions^1–3^. Likewise, gene tagging, e.g. with fluorescent protein reporters, has been used in functional genomics to study protein abundance^4^, localization^5^, turnover^6, 7^, or protein-protein interactions^8–10^.

Due to their gene-wise construction, producing arrayed clone collections is typically time-consuming and cost-intensive. For yeast, this has been partly addressed with the development of SWAT libraries in which a generic N- or C-terminal tag can be systematically replaced with the desired reporter for tagging any ORF in the genome^11, 12^. However, manipulation and screening of arrayed libraries remains dependent on special equipment to handle the strain collections and is confined to the genetic background of the yeast strain BY4741^13^ in which most of these libraries were constructed. Therefore, arrayed libraries cannot address current and future demands in functional genomics that embrace the systematic analysis of complex traits or the comparison of different strains or species^14^.

We imagine that a paradigm shift from arrayed to pooled library generation may offer a solution: Experimentation with pooled biological resources is already well established^15^ and the phenotype-to-genotype relationship can be inferred conveniently by genotyping phenotypically distinct subsets of pooled libraries using next-generation sequencing (NGS). To generate the pooled libraries rapidly and independent of their genetic background, an efficient strategy to introduce the genetic alterations is required. For example, RNA-programmable CRISPR-associated endonucleases have revolutionized the creation of pooled collections of gene activation and inactivation mutants in mammalian cells^16–18^ since thousands of CRISPR guide RNAs (gRNAs) can be produced by cost-effective microarray-based oligonucleotide synthesis. In bacteria and yeast, strategies that exploit homologous recombination have enabled multiplexed gene editing by delivering short repair templates on the same oligonucleotides as the gRNA^19–22^ with applications for phenotypic profiling of genomic sequence variations.

Because of high throughput, low cost, and broad host versatility, it is interesting to leverage these CRISPR-based methods beyond loss- or gain-of-function screens for the precise insertion of longer DNA constructs that deliver reporter molecules or tags to monitor the different cellular components encoded in the genome. Rapid access to such collections would synergize, e.g. with image-activated cell sorting^23^, and enable to use subcellular localization as a criterion for cell sorting.

To exert gene tagging in a pooled format, thousands of DNA constructs must be generated, each containing the reporter gene flanked with locus-specific homology arms and paired with a corresponding gRNA. However, parallel construction of thousands of such constructs is challenging.

Here, we describe ‘CRISPR-assisted tag library engineering’ (CASTLING) to create pooled collections of hundreds to thousands of yeast clones in a single reaction tube. All clones contain the same, large DNA construct (up to several kb in length) accurately inserted at a different, yet precisely specified chromosomal locus. CASTLING is compatible with microarray-based oligonucleotide synthesis since each insertion is specified by a single oligonucleotide only. Our method employs an intramolecular recombineering procedure that allows the conversion of oligonucleotide pools into pools of tagging cassettes.

In this proof-of-concept study, we establish CASTLING in the yeast *Saccharomyces cerevisiae* using gene tagging with fluorescent protein reporters as an example. We derive a set of rules to aid designing effective crRNAs for the CRISPR endonuclease Cas12a (formerly known as Cpf1)^24^ for C-terminal tagging of genes in yeast, and determine parameters to maximize tagging success in libraries of different sizes. We use a simple assay based on fluorescence-activated cell sorting (FACS) to demonstrate how CASTLING libraries can be used for proteome profiling and *ad hoc* characterization of previously uncharacterized proteins, and provide a targeted NGS method for the quantitative analysis of such pooled experiments.

## Results

### Gene tagging with self-integrating cassettes

The main component of CASTLING is a linear DNA construct that comprises multiple genetic elements: the ‘feature’ for genomic integration such as a fluorescent protein tag, a selection marker, a gene for a locus-specific Cas12a CRISPR RNA (crRNA) and flanking homology arms to direct the genomic insertion of the DNA fragment by homologous recombination (Figure 1a). We conceptually termed these DNA constructs ‘self-integrating cassettes’ (SICs).

**Figure 1.**
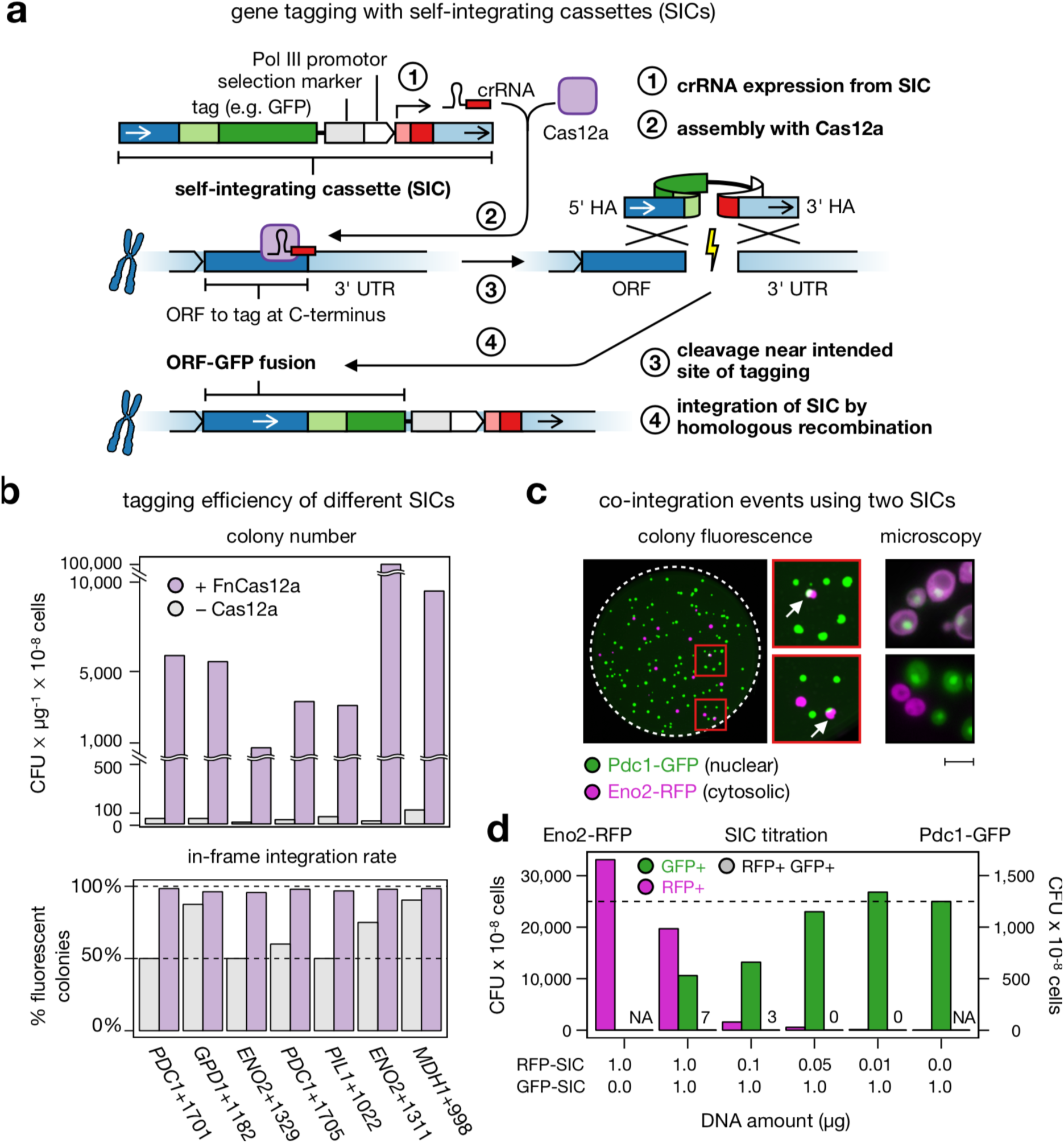
CRISPR-Cas12a-assisted single gene-tagging in yeast. (**a**) After transformation of the self-integrating cassette (SIC) into a cell, the crRNA expressed from the SIC directs a CRISPR-Cas12a endonuclease to the genomic target locus where the DNA double-strand is cleaved. The lesion is repaired by homologous recombination using the SIC as repair template so that an in-frame gene fusion is observed. (**b**) Efficiency of seven SICs of C-terminal tagging of highly expressed open-reading frames (ORFs) with a fluorescent protein reporter, in the absence (grey) or presence (purple) of FnCas12a. Colony-forming units (CFUs) per microgram of DNA and cells used for transformation, and integration fidelity by colony fluorescence are shown. (**c**) Co-integration events upon simultaneous transformation of two SICs directed against either *ENO2* or *PDC1*. Both SICs confer resistance to Geneticin (G-418), but contain different fluorescent protein tags. Colonies exhibiting green and red fluorescence (arrows) were streaked to identify true co-integrands. False-color fluorescence microscopy images show nuclear Pdc1-GFP in green and the cytosolic Eno2-RFP in magenta; scale bar 5 µm. (**d**) Titration of both SICs against each other (lower panel) with evaluation of GFP-tagged (GFP+), RFP-tagged (RFP+) or co-transformed (GFP+RFP+) colonies. Panels b–d: Source data are provided as a Source Data file.

We used Cas12a from *Francisella novicida U112* (FnCas12a), which is functional in yeast^25^, because the genomic target space of the Cas12a endonucleases is defined by A/T-rich protospacer-adjacent motifs (PAMs)^26–29^. This makes Cas12a endonucleases well suited for genetic engineering at transcriptional START and STOP sites in many organisms (Supplementary Figure 1).

To test the SIC strategy, we generated SICs for tagging several highly-expressed genes with a fluorescent protein reporter. After individual transformation of the SICs and marker selection, we obtained 100–1,000-times more colonies from hosts that had transiently expressed a Cas12a endonuclease as compared to a host that did not (Figure 1b). Also, the presence of a crRNA gene specific for the target locus of the SIC was required (Supplementary Figure 2), indicating that a functional crRNA transcribed from the linear DNA fragment promotes the integration of a SIC. Based on fluorescent colony counts, tagging fidelity had increased from 50–85% in the absence of Cas12a to 95–98% when recombination was stimulated by the action of Cas12a (Figure 1b).

We also tested Cas12a endonucleases from other species^24^, finding that Cas12a from *Acidaminococcus sp. BV3L6* (AsCas12a) showed similar activity as FnCas12a (Supplementary Figure 3a–c). However, we continued with FnCas12a since it offered a broader genomic target space in the yeast genome than AsCas12a (Supplementary Figure 4).

Because of the high efficiency of SIC integration, we worried that multiple loci could be tagged within the same cell when different SICs were transformed as pools. We therefore transformed a mixture of two SICs, one to tag *ENO2* with mCherry and the other one to tag *PDC1* with sfGFP. We detected only a few individual colonies where both genes were fluorescently tagged (Figure 1c), independent of the relative concentration of the two SICs used for transformation (Figure 1d). Therefore, tagging multiple loci in the same cell would rarely occur if more than one SIC was transformed simultaneously.

### Implementing CASTLING for pooled gene tagging

To produce many different SICs in a pooled format using microarray-synthesized oligonucleotides, all gene-specific elements of a SIC, i.e. the crRNA sequence and both homology arms, must be contained in a single oligonucleotide—one for each target locus (Figure 2a). In turn, this demands a strategy to convert these oligonucleotides in bulk into the corresponding SICs.

**Figure 2.**
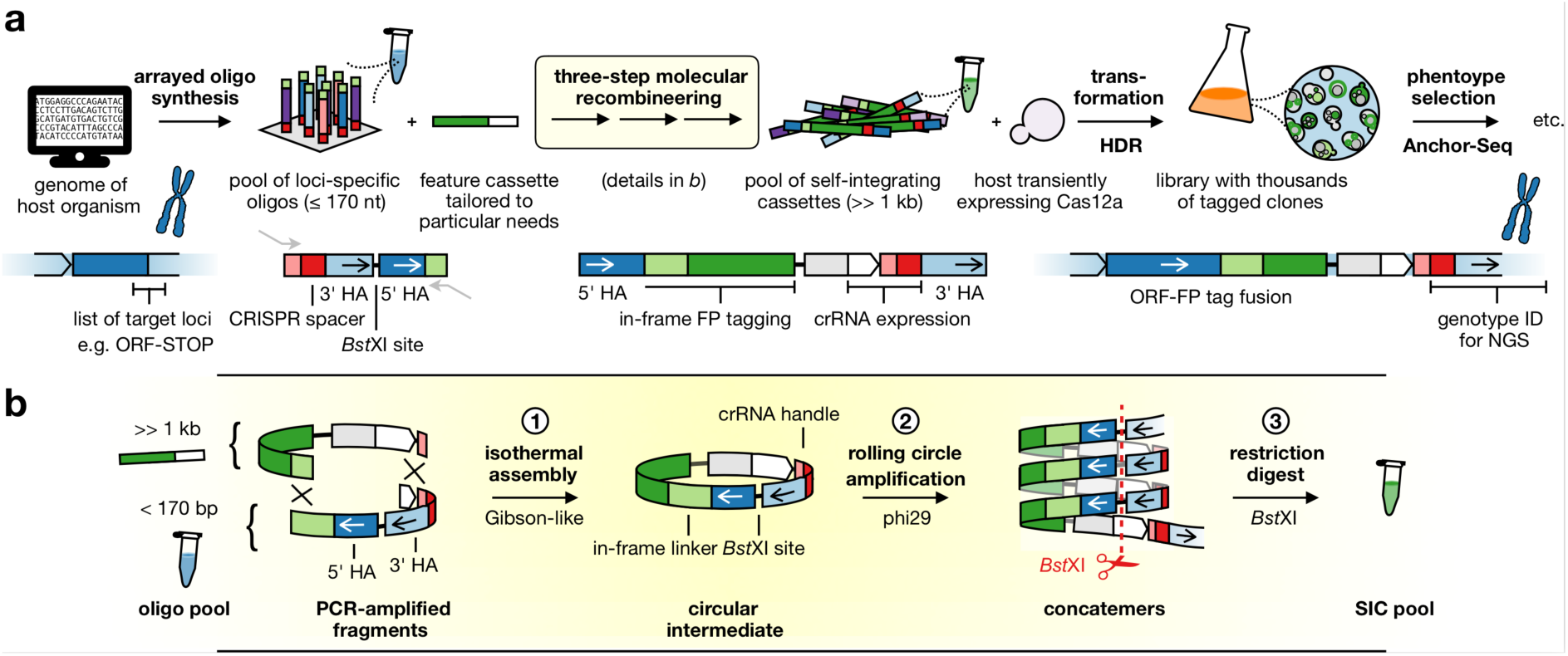
CASTLING in a nutshell. (**a**) For each target locus, a DNA oligonucleotide with site-specific homology arms (HA) and a CRISPR spacer encoding a target-specific crRNA is designed and synthesized as part of an oligonucleotide array. The resulting oligonucleotide pool is recombineered with a custom-tailored feature cassette into a pool of self-integrating cassettes (SICs). This results in a clone collection (library) that can be subjected to phenotypic screening and genotyping, e.g. using Anchor-Seq^12^. (**b**) The three-step recombineering procedure for SIC pool generation; details in the main text and Methods.

We implemented a three-step molecular recombineering procedure for this conversion that is executed *in vitro* (Figure 2b, Supplementary Figure 5a–e). Its central intermediate is a circular DNA species formed by the oligonucleotides and a feature cassette. The feature cassette provides all the generic elements of the SIC, i.e. the tag (e.g. GFP), the selection marker and an RNA polymerase III (Pol III) promoter to express the crRNA. The circular intermediates are then amplified by rolling circle amplification (RCA) instead of PCR to avoid the formation of chimeras containing non-matching homology arms. The individual SICs are finally released by cleaving the DNA concatemer using a restriction site in between both homology arms.

To accommodate all gene-specific elements on a single oligonucleotide, it was critical to use a Cas12a endonuclease because its crRNA consists of a comparably short direct repeat sequence (∼20 nt) that *precedes* each target-specific CRISPR spacer (∼20 nt; Supplementary Figure 5f). This arrangement allows the Pol III promoter, which drives crRNA expression, to remain part of the feature cassette, while the short Pol III terminator^30^ can be included in the oligonucleotide itself. This design leaves enough space for homology arms of sufficient length for homologous recombination (>28 bp) ^31^. Adding up all the sequences, each oligonucleotide (160–170 nt) is within the length-limits for commercial microarray-based synthesis.

To select CRISPR targets near the desired chromosomal insertion points and to assist the design of the oligonucleotide sequences for microarray synthesis (Supplementary Figure 6a–d), we wrote the software tool castR (https://github.com/knoplab/castR/tree/v1.0[https://github.com/knoplab/castR/tree/v1.0]). For use with small genomes, castR is available online (http://schapb.zmbh.uni-heidelberg.de/castR/[http://schapb.zmbh.uni-heidelberg.de/castR/]).

### Using CASTLING to generate a GFP library of nuclear proteins

To test CASTLING, we sought to create a small library covering a set of proteins with known localization^32^. We chose 215 nuclear proteins whose localization had been validated in different genome-wide datasets^12, 33^. We designed 1,577 oligonucleotides covering all suitable PAM sites within 30 bp around the C-termini of the selected ORFs, yielding 7 oligonucleotides per gene on average. We purchased this oligonucleotide pool three times from different suppliers, one pool from supplier A (pool A) and two pools from supplier B (pool B1 and B2; Figure 3a, Methods). The amount of starting material for PCR to amplify each pool was adjusted to obtain a product within ∼20 cycles. We observed that pool A required about 200-fold more starting material than pool B1 or B2 (Figure 3a). After recombineering with a feature cassette comprising the bright green fluorescent protein reporter mNeonGreen^34^, we generated four different libraries in technical duplicates of 30,000–95,000 clones each (Figure 3a, Supplementary Table 1).

**Figure 3.**
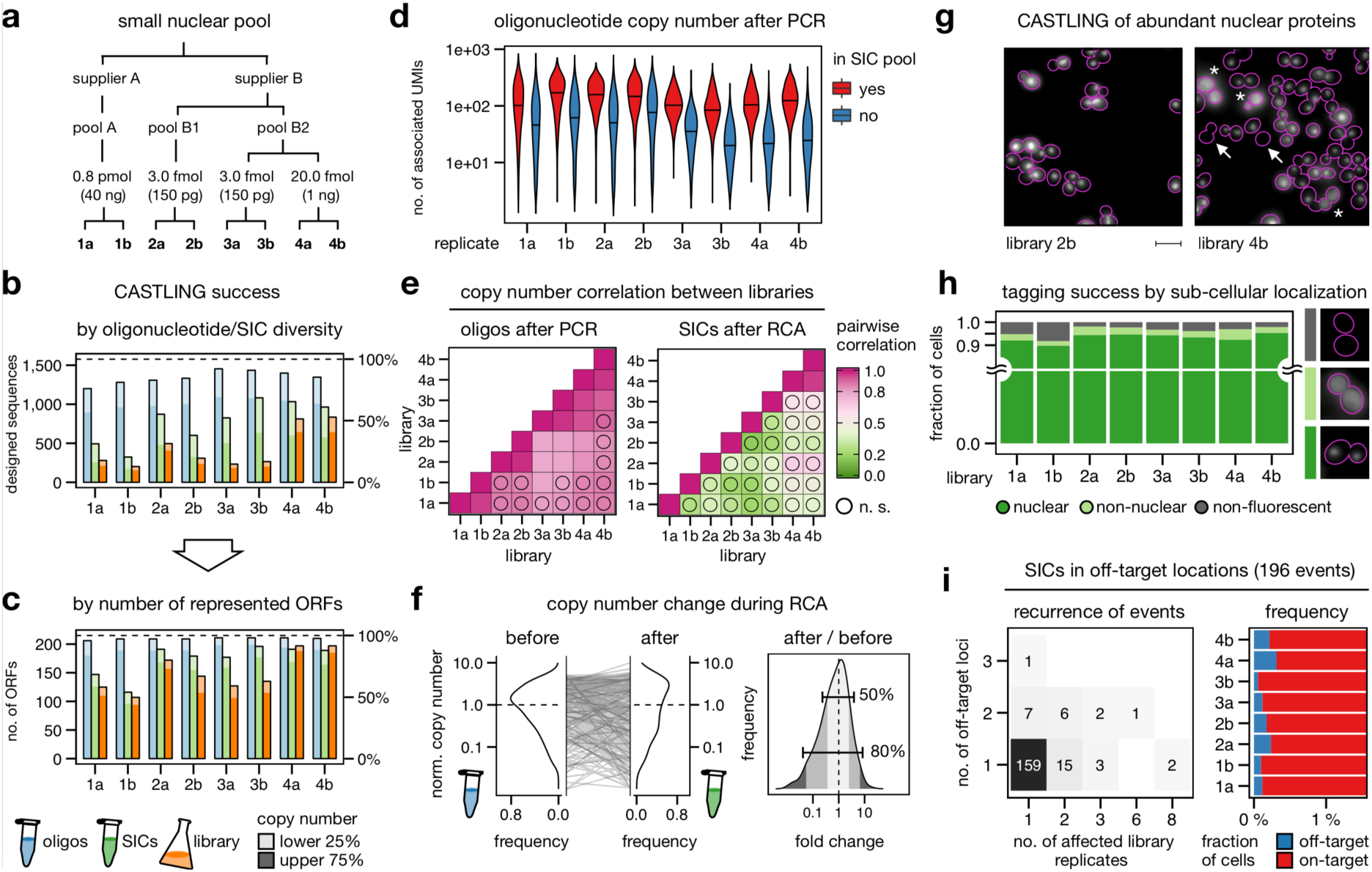
CASTLING for tagging 215 nuclear proteins with a green fluorescent protein. (**a**) Three oligonucleotide pools of the same design (1,577 sequences, Supplementary Table 1) were used to create four tag libraries by CASTLING in duplicate sampling the indicated amount of starting material for PCR. (**b**) Detected oligonucleotide sequences of the design after PCR amplification (blue), self-integrating cassette (SIC) assembly (green) and in the final library (orange); oligonucleotides with copy number estimates (unique UMI counts) in the lowest quartile (lower 25%) are shown in light shade. (**c**) Same as (b), but evaluated in terms of open reading frames (ORFs) represented by the oligonucleotides or SICs. (**d**) Copy number of PCR amplicons recovered (red) or lost (blue) after recombineering; black horizontal lines indicate median UMI counts. (**e**) Pearson’s pairwise correlation of oligonucleotide or SIC copy number between replicates after PCR or rolling-circle amplification (RCA) respectively; n.s., not significant (p > 0.05). (**f**) Kernel density estimates of copy number in replicate 1a as normalized to the median copy number observed in the oligonucleotide pool (before recombineering) and after recombineering into the SIC pool (left panel); the distribution of fold-changes (right panel) highlights two frequency ranges: [0.1–0.9], i.e. 80% of SICs, and [0.25–0.75], i.e. 50% of SICs. (**g**) Representative fluorescence microscopy images of cells displaying nuclear, diffuse non-nuclear (asterisks), or no mNeonGreen fluorescence (arrows); scale bar 5 µm. (**h**) Quantification of fluorescence localization in >1,000 cells in each replicate. (**i**) Recurrence of off-target events as revealed by Anchor-Seq across all library replicates and all genomic loci (left panel); the fraction of cells with SICs integrated at off-target sites (blue) within each clone population (red) is shown (right panel, axis trimmed). Panels b–i: Source data are provided as a Source Data file.

We used NGS in combination with unique molecular identifiers (UMIs)^35^ to quantitatively analyze the entire procedure at three stages: After PCR amplification of the oligonucleotide pool, after SIC amplification (Supplementary Figure 7a), and after yeast library construction. To characterize the yeast libraries, we adapted the targeted NGS method Anchor-Seq^12^ with UMIs to analyze the CRISPR spacers of the inserted SICs along with the genomic sequence adjacent to the insertion site in all clones of the libraries (Supplementary Figure 7b).

Overall, the represented oligonucleotide diversity gradually decreased during recombineering (Figure 3b). The best performance was observed in one duplicate generated from pool B2 that used a high amount of starting material (libraries 4a and 4b), preserving more than 70% of the originally amplified oligonucleotides in the SIC pool and more than 60% of the oligonucleotide diversity in the yeast libraries (Figure 3b). This loss in complexity was alleviated by the fact that multiple oligonucleotides were included per gene and we observed that more than 90% of the targeted genes were tagged in library 4a and 4b (Figure 3c). We noticed that low abundant oligonucleotides after PCR amplification were prone to depletion during SIC preparation, accounting for the observed loss in sequence diversity (Figure 3d). Across all preparations, copy numbers of individual oligonucleotides were highly correlated between duplicates after PCR (Pearson correlation >0.96), but less between synthesis replicates (0.78–0.90), and least for oligonucleotide pools obtained from different suppliers (Figure 3e). After recombineering and RCA, no significant correlation of SIC copy numbers was observed except for libraries 4a and 4b. A more detailed analysis indicated that 50% of the sequences exhibited a copy number change >2-fold during RCA (Figure 3f), which could explain the loss of correlation between replicates after RCA. Taken together, these analyses identified the quality and amount of starting material and its recovery during recombineering as critical factors to preserve library diversity. Nevertheless, for a small library of 215 genes, CASTLING enabled tagging most of the selected genes within one library preparation.

Next, we quantified tagging fidelity by fluorescence microscopy, which was possible because we had selected genes encoding proteins with validated nuclear localization: 90–95% of the cells had a nuclear localized mNeonGreen signal in all libraries (Figure 3g–h). The remainder of the cells showed either no fluorescence (2–8%) or a fluorescence signal elsewhere (0–4%), usually in the cytoplasm with one exception (see below). So, nearly all genes must have been tagged in the correct reading frame.

For the clones with no fluorescence signal, we suspected either frame-shift mutations in the polypeptide linker (due to faulty oligonucleotides) or in the fluorescent protein reporter (due to limited fidelity of DNA polymerases), or off-target integration of the SIC. Sequencing of several insertion junctions of dark clones revealed small deletions of one or more nucleotides in the 5’-homology arms that direct the SICs to the 3’-ends of the ORFs. Therefore, the majority of dark clones appeared to contain correctly targeted SICs in which mNeonGreen was not in frame due to errors in the sequences derived from the oligonucleotides.

Next, we generated library-wide Anchor-Seq data encompassing the crRNA sequences and the 3’-insertion junctions. This identified 280 instances in which the crRNA sequence and the genomic insertion site did not match. These off-target insertions corresponded to less than 0.2% of the clones. Most of them were single occurrences associated with 196 different SICs in total. Only 37 SICs showed off-target insertion at various genomic loci or in more than one library replicate (Figure 3i). It remains however unclear, which of these insertions were caused by Cas12a-mediated cleavage at an off-target site and which were spontaneous chromosomal insertions.

In addition to these events, we observed fluorescence signals at unexpected subcellular localizations. For example, 2% of the cells in library 2b displayed fluorescence at the spindle-pole body, which we attributed, based on Anchor-Seq, to a *TEM1*-mNeonGreen gene fusion. Indeed, on average 1.6% of all cells across all libraries had integrated SICs originally designed for another experiment in this study, which must have entered SIC or library preparation as a result of contamination.

Together, these experiments demonstrate that in a pooled experiment CASTLING allows for highly efficient tagging of hundreds of genes with low levels of off-target insertion.

### Parameters affecting tagging success on a genome-wide scale

Simultaneously with the small pool of nuclear proteins, we designed an oligonucleotide pool for C-terminal tagging of the yeast proteome. For crRNA design, we first retrieved a set of more than 34,000 candidate CRISPR targets by castR using TTV (V = A, C, or G) and TYN (Y = C or T; N = any nucleobase) as protospacer-adjacent motifs (PAMs). Next, we removed sequences that contained thymidine runs longer than five nucleotides, since they may prematurely terminate Pol III transcription^30^. Subsequently, we filtered out crRNA targets with a high off-target estimate and removed most, but not all target sequences that are not destroyed after insertion of the SICs (Supplementary Note 1). From the remainder, we chose randomly 12,472 sequences (limited by the chosen microarray) that covered 5,664 of 6,681 (85%) of the annotated ORFs in *S. cerevisiae*^36^. Although the number of oligonucleotides per gene was lower as compared to the nuclear pool, the high number of genomic targets should allow identifying parameters that would influence tagging success and clone representation in such large-scale experiments.

After PCR and SIC pool generation, we sequenced the PCR amplicons and one SIC pool. We analyzed the sequencing data implementing a de-noising strategy to discriminate errors introduced during NGS from errors in the templates^37^. This revealed that the PCR product contained 57% of the designed oligonucleotides, but only 31% of the designed sequences were represented by at least one error-free amplicon. Similarly, 51% of all designed sequences were detected in this SIC pool, but only 25% were error-free (Figure 4a). Due to redundancy, the error-free SICs in this pool still covered 45% of the 5,664 ORFs.

**Figure 4.**
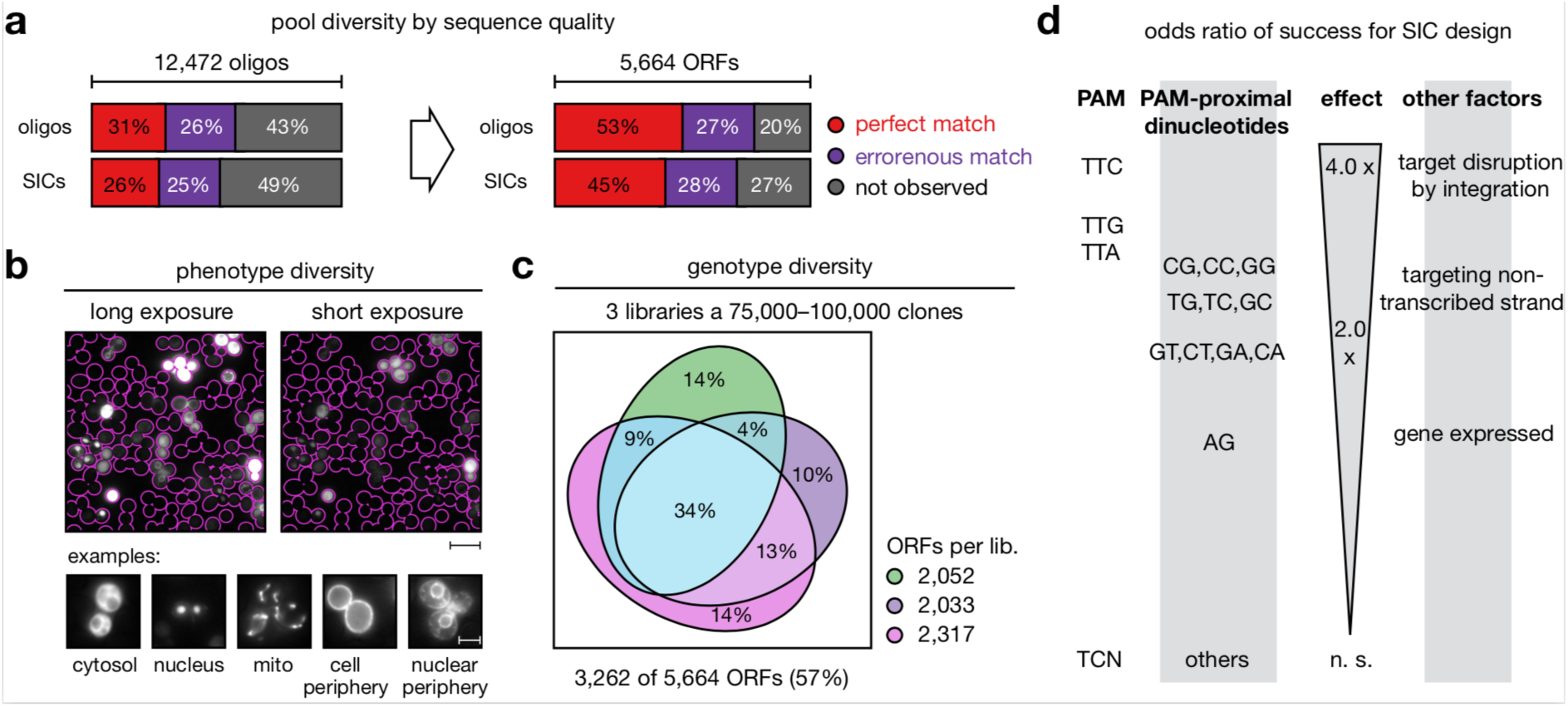
Identification of factors influencing clone representation in CASTLING. (**a**) Sequence quality of an oligonucleotide pool (oligonucleotide pool C, Supplementary Table 2) after PCR amplification and SIC assembly. Following de-noising of NGS artifacts, molecules that aligned with any of the 12,472 designed oligonucleotides were classified error-free, erroneous or absent at the respective stage (left panel). The genotype space (designed: 5,664 ORFs) covered by each class (right panel). (**b**) Representative fluorescence microscopy images of a pooled tag library (derived from oligonucleotide pool C); scale bar 20 µm (overview), 5 µm (details). (**c**) Genotype diversity within three independent library preparations (libraries #1.1, #1.2 and #1.3, Supplementary Table 2) generated from the same oligonucleotide pool; all libraries combined tagged 3,262 different ORFs. (**d**) Summary of parameters significantly (Fisher’s exact test, p < 0.05) increasing the likeliness of tagging success beyond SIC abundance (details in Supplementary Figure 7a–b). Panels a–c: Source data are provided as a Source Data file.

To explore how many genes could be tagged with this oligonucleotide pool, we repeated PCR and SIC assembly three times. Following transformation in yeast, this resulted in three independent libraries of 75,000–100,000 clones each. Inspection of the cells by fluorescence microscopy revealed localization across a broad range of subcellular compartments (Figure 4b). By Anchor-Seq, we detected a total of 3,262 different ORFs (58% of all targeted ORFs), of which 1,127 ORFs (20%) were shared across all replicates (Figure 4c, Supplementary Table 2).

The acquired data allowed us to identify factors that might have impeded efficient genomic integration of a SIC. First, the likelihood of tagging success was 3- to 4-fold decreased when the crRNA target sequence was not disrupted by the inserted SIC, i.e. when recurrent cleavage of the locus was possible. Neither nucleosome occupancy of the PAM, nor of the target sequence itself had a statistically significant impact on the tagging success in this library. However, the choice of the PAM (TTC>TTG>TTA >> TYN) and the first two PAM-proximal nucleotides (CG, CC, GG) increased the chances of target integration 2- to 3-fold each (Figure 4d, Supplementary Figure 8a–b). Interestingly, it seemed advantageous to target genes on their non-transcribed strand by Cas12a. Despite the limited success to create a genome-wide library at first trial, we anticipated that these parameters could help to improve tagging success for CASTLING in yeast.

### Using CASTLING to construct complex pooled yeast libraries

To further investigate the creation of genome-wide pooled libraries with CASTLING, we designed a new microarray for tagging 5,940 ORFs. Applying these rules for each ORF, we selected 17,691 target sites near the STOP codon and filled up the remaining positions on a 27,000-well array. We generated three libraries in total using two different strategies to investigate the minimal effort that would be required for creating a large library with CASTLING. First, we pooled SICs from 30 RCAs and generated a large library of 704,000 clones (LibA), and a small library of 44,000 clones (LibB). Second, we constructed a third library of 116,000 clones (LibC) using a SIC pool made from two RCAs of the same oligonucleotide pool (Figure 5a). To quantify genotype composition in each of the different libraries, we again used Anchor-Seq at the crRNA junction. Altogether, the three libraries contained tagged alleles of 76% of all the targeted ORFs with an overlap of 43% between the three libraries (Figure 5b–c). The largest library, LibA, contained the most tagged ORFs (3,801 ORFs), corresponding to 64% of the design. Interestingly, the much smaller library LibB with 44,000 clones already contained 80% of these genotypes. LibB and LibC each covered ∼50% of the desired ORFs, sharing 2,038 ORFs. In practical terms, this implied that about one third of the intended genes could be reliably and reproducibly tagged with minimal effort by recovering 40,000–120,000 clones only.

**Figure 5.**
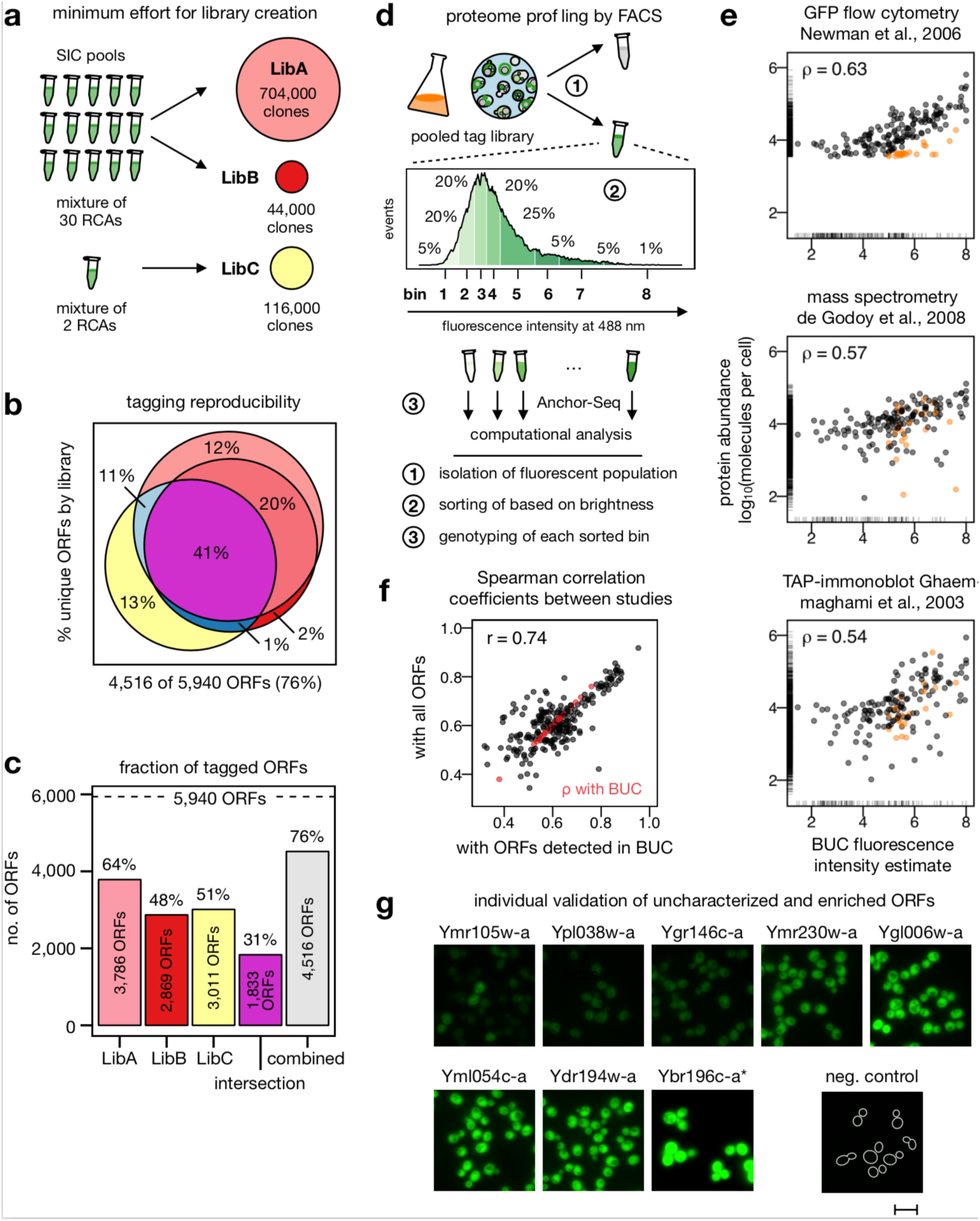
Creating and screening large CASTLING libraries. (**a**) Three libraries with different numbers of collected clones were generated from SIC pools combining either 2 or 30 recombineering reactions to investigate the minimum effort for a proteome-wide (design: 5,940 ORFs, oligonucleotide pool D, Supplementary Table 2) CASTLING library (details in Methods). (**b**) Venn diagram of genotypes recovered in each of the three libraries; all libraries combined tagged 4,516 different ORFs. (**c**) Genotype diversity in each of the three libraries, shared between them, or after their combination. (**d**) Proteome profiling by fluorescence intensity of a non-exhaustive mNeonGreen tag library (library #1.1, Figure 4c, Supplementary Table 2) using FACS. After enriching the fluorescent sub-population of the library and determining the fold-enrichment of each genotype by NGS, this sub-population was sorted into eight bins according to fluorescent intensity. Analysis of each bin by Anchor-Seq and on-site nanopore sequencing allowed the assignment of an expected protein abundance for each genotype. (**e**) Pairwise comparisons between fluorescence intensity estimates calculated from genotype distribution across all bins (Methods, Equation 2; this study denoted as BUC) and protein abundances reported by selected genome-scale experiments^4, 39, 40^ normalized to molecules per cell^38^. Outliers (orange) were determined based on the comparison to a GFP tag flow cytometry study^39^. Spearman correlation coefficients (ρ) are given. Marginal lines indicate abundance estimates only present in the respective study but missing in the other. (**f**) Comparison of Spearman correlation coefficients between studies either considering their overlap in detected ORFs or only the overlap with the 435 ORFs we could detect in this experiment. A Pearson correlation coefficient (r) is given. (**g**) Eight genes that had not been characterized in other genome-scale experiments^38^ were tagged individually to verify whether fluorescence intensity corresponded with their predicted characterization by FACS. Same exposure time for all fluorescent microscopy images except for Ybr196c-a which was imaged at 10% excitation; scale bar 10 µm. Panels b, c and e: Source data are provided as a Source Data file.

We validated the rule set used for oligonucleotide design by comparing SICs with approximately equal copy number in the SIC pool (Supplementary Figure 9a–b).

Functional studies that use pooled libraries fundamentally depend on enrichment procedures to physically separate cells based on the information provided by the reporter. When fluorescent protein fusions to endogenous proteins are used, high-resolution fluorescence microscopy would be the method of choice, as this would enable scoring and subsequent cell sorting based on very complex but highly informative phenotypes. The necessary technology is currently under development^23^. To demonstrate that CASTLING libraries can be used for screening, we reverted to FACS which permits sorting based on fluorescence intensity.

Starting from a library containing 2,052 mNeonGreen-tagged ORFs (Figure 4c), we first sorted cells for which fluorescence could be detected by FACS. Anchor-Seq revealed that in comparison to the starting library, this cell population contained 848 genotypes while 732 genotypes were depleted. Therefore, we estimated that 35% of the mNeonGreen-tagged genes could be profiled based on fluorescence intensity in our pooled study, which agrees with a meta-analysis on yeast protein abundance^38^ that reported abundance estimates for 1,404 proteins characterized by flow cytometric fluorescence measurements^39^, i.e. 34% of 4,159 ORFs tagged in the C-GFP library^32^.

To determine the fluorescence intensity of individual proteins in the fluorescence-enriched fraction, we sorted the cells into 8 fractions of increasing fluorescence intensity Next, we analyzed the genotype distribution within the bins using Anchor-Seq. We sequenced the amplified insertion junctions using MinION nanopore sequencing. This method allows a more rapid profiling workflow but provides a lower sequencing depth as compared to Illumina dye sequencing, which we usually used to characterize CASTLING libraries. We obtained 18,638 informative reads, which enabled us to determine the relative enrichment in the individual bins for 435 (50%) of the 848 tagged proteins. These estimates correlated well with the abundance estimates from the flow cytometry study by Newman *et al.*^39^ (Spearman correlation coefficient >0.63; Figure 5e) and were comparable with different protein abundance data sets consolidated by Ho *et al.*^38^ (Supplementary Figure 10).

To estimate whether our low-depth showcase experiment can be considered representative for larger-scale CASTLING-based experiments, we quantified the dependence of correlation coefficients on the number of compared genes and found that the coefficients of correlation obtained from the complete data sets^38^ or an analysis limited to the 435 tagged genes that we had detected, correlated well with each other (Pearson correlation coefficient 0.74; Figure 5f), indicating a predictive value of our low-depth experiment.

We found that 23 (13%) of 175 tagged genes yielded clearly detectable fluorescence signals in our study, but were not detected by Newman *et al.*^39^ (Figure 5e, orange points). Since these proteins were also detected by complementary approaches such as mass spectrometry^40^ or immuno-blotting^4^, we assumed that these ‘false positives’ resulted from false negative clones of the C-GFP library (Figure 5e). Using independently generated clones based on a different gene tagging strategy^12^, we validated the expression of most of these genes when tagged with mNeonGreen, including 8 proteins that were not covered in the C-GFP library and neither characterized in any other study analyzed by Ho *et al.*^38^ (Figure 5g, Supplementary Table 3).

Together, these results highlight the use of CASTLING libraries as a rapid venue for phenotypic profiling and screening experiments when combined with Anchor-Seq to analyze the clone distribution across sub-populations isolated from such libraries.

## Discussion

We developed CASTLING to enable the rapid creation of pooled libraries of clones with large chromosomal insertions such as fluorescent protein tags.

Typically, libraries in yeast have been constructed gene-wise in an arrayed format using PCR-targeting^41^. Based on our own experience^12, 42^, the construction of arrayed libraries depends on special equipment for parallelization of the procedures, it requires a (costly) resource of arrayed primers for PCR tagging, and handling thousands of strains keeps multiple researchers occupied for several months.

In contrast, fewer resources must be committed to create a library by CASTLING. All the necessary oligonucleotides can be obtained from microarrays, which are about two orders of magnitude more cost-effective than a genome-wide set of conventional solid-phase-synthesized oligonucleotides. Once an established oligonucleotide pool is available, it can be reused to construct a variety of SIC pools containing different features, i.e. tags or selection markers. The construction of SIC pools is rapid and can be completed within 1–2 days since the CASTLING workflow avoids preparatory sub-cloning into a plasmid library, which is commonly used in other multiplexed gene-editing approaches^19–21, 43^. Transformation and growth of the yeast clones take another 2–3 days, followed by recovery and analysis of the library. This makes library preparation by CASTLING very efficient and therefore it is possible to create a new library for each strain background or mutant of interest. Classical libraries in contrast, are confined to the background they were made in and require genetic crossing to introduce a mutant, which depends on strains specifically constructed for these procedures^44^.

In addition to the versatility and flexibility of library creation, tagging fidelity by CASTLING is 90% or higher, exceeding the fidelity observed in conventional gene tagging by PCR-targeting, where routinely 50–85% of the obtained clones are correct. It may be worth mentioning that elimination of the false clones during the construction of classical arrayed libraries remains one of the most laborious steps. With CASTLING, false clones cannot obstruct the correct interpretation of a screening because with Anchor-Seq all genotypes can be quantified that are present at the beginning of an experiment as well as their respective enrichment or depletion after phenotypic selection. This allows excluding erroneous genotypes while completing the analysis, which is typically not possible in other multiplexed CRISPR-based gene editing approaches that rely on indirect measures for genotype determination (e.g. sequencing the ectopic crRNA plasmids).

A potential down-side of CASTLING and many other pooled library approaches lies within the initial indeterminacy of the exact library composition: Each transformation will yield pools with not exactly the same composition. Currently, genotype coverage with CASTLING can exceed 90% when relatively small libraries with hundreds of genes are created and reproducibly reached 50% for libraries with thousands of genes using less than 10-fold oversampling (44,000 clones over 5,940 ORFs). We have identified that SICs for which the CRISPR target site would be destroyed after integration, or SICs that had a GC-rich crRNA in its PAM-proximal dinucleotides, yielded higher clone numbers as compared to SICs lacking those features (Supplementary Figure 9).

The identified parameters increased the likelihood of tagging success, but they might also reduce the number of clones for ORFs for which only less efficient SICs could be designed. In this case, additional oversampling would be required. Along this line, a better strategy to increase coverage might be to use successive rounds of CASTLING involving each time a new microarray to target the remainder of genes. The first array would target those genes that can be reproducibly tagged in all trials (Figure 5b–c), while subsequent arrays would incrementally complete the library with almost proportional scaling efforts in terms of clones to be collected. Probably, it would require 2–4 rounds of CASTLING with a total of 60,000–120,000 clones to tag >60–90% of all 5,500–6,000 genes in yeast. This would exceed available genome-wide tagging collection, e.g. the C-GFP collection^32^ with 4,159 ORFs (Thermo Fisher), the TAP-tag collection^4^ with 4,247 ORFs (Dharmacon) or our tandem fluorescent timer collection with 4,081 ORFs^42^. Importantly, such an optimization might be necessary only once; Afterwards, all oligonucleotide pools could be used in parallel to generate a nearly complete library. This approach might also yield optimized rule sets to guide the development of CASTLING for a different species.

A major factor that decreased tagging success seemed to be oligonucleotide quality. CASTLING requires long oligonucleotides >100 bp. Even very small error rates and almost perfect coupling efficiencies during oligonucleotide synthesis will give rise to pools that only contain a minor fraction of full-length error-free oligonucleotides. Furthermore, we observed that the same sequences synthesized in different batches gave rise to pools with different performance (Pool B1 and B2). We have sequenced and thoroughly analyzed one of the oligonucleotide pools for large library creation. Only a fraction of the designed sequences was represented by perfect full-length oligonucleotides. Most frequently we observe deletions and SNPs in the oligonucleotide sequences. SNPs seem to be more frequent at the 3’ end of the oligonucleotide (which is synthesized first), whereas deletions become more frequent towards the 5’ end of the oligonucleotide (which is synthesized last). Indeed, error-free synthesis of long oligonucleotides remains challenging^45, 46^. To increase the chance of representing each target locus by a perfect oligonucleotide, it might be beneficial to use as many different oligonucleotides per gene as possible or to include multiple redundant sequences.

It is important to stress that faulty oligonucleotides do *not* necessarily impact the fidelity of the tagging because the *in vitro* recombineering steps and the *in vivo* recombination^47^ all select against faulty oligonucleotides. Also, errors in the crRNA will most likely render it inactive. Consequently, only a few oligonucleotides that end up in the genome are associated with frameshift errors that impair the expression of the tag. This is impressively demonstrated with the nuclear protein libraries that were prepared with three different oligonucleotide pools, all of which showing >90% in-frame tagging rates (Figure 3h). This results in intrinsic quality control during CASTLING yielding correctly tagged genes in the majority.

### Prospective applications of CASTLING

In combination, CASTLING and quantitative Anchor-Seq enable the rapid creation and analysis of pooled libraries with tagged genes. Since each reaction tube contains an entire library, the pooled format is able to address much broader, comparative questions, including different genetic backgrounds and/or environmental conditions.

CASTLING is a method for gene tagging, and the type of screen that can be performed with such libraries entirely depends on the used tag. Therefore, it is up to the creativity of the researcher to develop a screening procedure to convert the information provided by the tags into information about the biological question in mind. Importantly, a screening procedure requires physical fractionation of the library into sub-pools based on a suitable phenotypic read-out, for example using tags that enable the coupling of a protein behavior such as protein localization^48^ or protein-protein interactions^10^ with a growth phenotype.

In our opinion, fluorescent protein reporters constitute a particularly attractive group of tags as they provide visual insights into the cellular organization and dynamics, changes of which are associated with many disturbances of biological processes. Our simple FACS enrichment experiment (Figure 5d–g) can serve but as proof of principle in this regard as current flow cytometry-based cell sorters cannot resolve complex cellular phenotypes such as the subcellular localization of proteins^49^. We think that for methods such as the recently developed image-activated cell sorting^23^, CASTLING can enable a variety of entirely new experimental designs and analyses, ranging from functional genomics to biomedical research, paving the way to a new paradigm of shot-gun cell biology.

Beyond yeast, CASTLING could be adapted for other organisms able to repair DNA lesions by homologous recombination, including bacteria, fungi, flies and worms, and potentially also in plants and mammalian cells. First evidence that this is the case is provided in Fueller et al.^50^ where we show that an adapted SIC strategy can be used for efficient endogenous tagging of genes in mammalian cells. We have preliminary data suggesting that CASTLING also works in mammalian cells, although the size of the library that can be generated with it is currently unclear.

In summary, our work shows that CASTLING libraries and quantitative genotype analysis using Anchor-Seq seamlessly integrate into existing (and upcoming) high-throughput cell sorting instrumentation to enable functional analyses of pooled resources. This outlines new avenues for the investigation of complex cellular processes in direct competition with strategies based on arrayed library resources.

*Please note*: Inadequate adoption of CASTLING can unwittingly generate clones qualified to initiate a gene-drive upon sexual reproduction^51, 52^. This can be easily prevented (Supplementary Note 2).

## Methods

### Yeast strains and plasmids

All strains were derived from ESM356-1 (*Saccharomyces cerevisiae* S288C, *MAT***a** *ura3*-52 *leu2*Δ1 *his3*Δ200 *trp1*Δ63, which is a spore from strain FY1679^1, 2^) and are listed in Supplementary Table 4. Plasmids are listed in Supplementary Table 5. Human codon-optimized Cas12a (formerly Cpf1) family proteins^3^ of *Francisella novicida U112* (FnCas12a), *Lachnospiriceae bacterium ND2006* (LbCas12a), *Acidaminococcus sp. BV3L6* (AsCas12a), and *Moraxella bovoculi 237* (MbCas12a) were expressed using the galactose-inducible *GAL1* promoter^4^ from plasmids integrated into the *ura3*-52 locus (pMaM486, pMaM487, pMaM488, pMaM489).

### Cell lysis and Western blot detection of HA tagged proteins

Proteins were extracted as described^5^ and resolved on Tris-glycine buffered 10% (v/v) polyacrylamide gels by electrophoresis at 200 V for 90 min, transferred onto a nitrocellulose membrane by wet blotting (12 mM Tris, 96 mM glycine, 20% (v/v) methanol) at 25V for 120 min, blocked with 10% (w/v) milk powder in blotting buffer (20 mM Tris, 150 mM NaCl, 0.1% (w/v) Tween 20) and the proteins of interest detected with monoclonal mouse anti-Pgk1 (R & D Systems, Fisher Scientific) and monoclonal mouse anti-HA (12CA5, Sigma-Aldrich) antibodies bound in 5 % (w/v) milk powder in blotting buffer at 4 °C overnight. The surplus of unbound primary antibody was washed away and the secondary HRP-coupled antibody applied in 5 % (w/v) milk powder in blotting buffer at room temperature.

### CASTLING library design

To facilitate oligonucleotide design, an *R* package is available from our repository (https://github.com/knoplab/castR/tree/v1.0[https://github.com/knoplab/castR/tree/v1.0]) that ships along with a GUI. For small genomes, the GUI can be accessed online (http://schapb.zmbh.uni-heidelberg.de/users/knoplab/castR/[http://schapb.zmbh.uni-heidelberg.de/users/knoplab/castR/]). The principles used for oligonucleotide design are described in Supplementary Note 1.

Oligonucleotide sequences used for microarray synthesis of oligopools in this study are given in Supplementary Data 1 (for arrays used in Figure 3), Supplementary Data 2 (for the array used in Figure 4), and Supplementary Data 3 (for the array used in Figure 5).

### Generating self-integrating cassettes for individual genes

Individual SICs were generated by PCR using a corresponding plasmid template (Supplementary Table 5) and using primers (Supplementary Table 6) that introduced the required 5’ and 3’ homology arms along with a locus-specific crRNA spacer. Cycling conditions for VELOCITY DNA polymerase-based amplification (Bioline) were 97 °C for 3 min, followed by 30 cycles of 97 °C (30 sec), 63 °C (30 sec), 72 °C (2 min 30 sec) and a final 72 °C (5 min) extension hold. The reactions were column-purified and adjusted to equal SIC concentration before yeast cell transformation.

### Amplifying oligonucleotide pools and feature cassettes

The oligonucleotide pools used in this study (Supplementary Table 7) were synthesized by either CustomArray Inc. (pool A and C), Twist Bioscience (pool B1 and B2), or Agilent Technologies (pool D), and reconstituted in TE in case they arrived lyophilized. Pool dilution and annealing temperature were optimized in each case to yield a uniform product of the expected length (Supplementary Figure 5a, Supplementary Table 1–2). In this study, pool C was diluted 1,000-fold and 1.5 fmol were amplified using VELOCITY DNA polymerase (Bioline) with forward primer pool-FP1 and reverse primer pool-RP2 using the following PCR conditions: 97 °C for 3 min, followed by 20 cycles of 97 °C (30 sec), 58 °C (30 sec), 72 °C (20 sec), and a final 72 °C (5 min) extension hold. To keep library member representation as uniform as possible, using more input material and higher annealing temperatures is desirable, as this will usually require fewer PCR cycles for amplification of the full-length synthesis product. All other pools were designed to allow for amplification in 15 cycles using Herculase II DNA polymerase (Agilent Technologies) with forward primer pool-FP2 (or pool-FP3, as indicated) and reverse primers pool-RP2 (or pool-RP3). Cycling conditions were: 95 °C for 2 min, followed by 6 cycles of 95 °C (20 sec), touch-down from 67 °C (20 sec, ΔT = –1 °C per cycle), 75 °C (30 sec), then 9 cycles of 95 °C (20 sec), 67 °C (20 sec), 72 °C (30 sec), and a final 72 °C (5 min) extension hold. Primers and truncated oligonucleotides (< 75 bp) were removed using NucleoSpin Gel and PCR clean-up columns (MACHERY-NAGEL GmbH & Co. KG). Feature cassettes were amplified by PCR using cognate cassette-FP and cassette-RP and any compatible plasmid template (50 ng, Supplementary Table 5) under the following conditions: 97 °C for 3 min, followed by 30 cycles of 97 °C (30 sec), 63 °C (30 sec), 72 °C (2 min 30 sec) and a final 72 °C (5 min) extension hold. The reaction was treated with *Dpn*I (New England Biolabs) *in situ* and cleaned-up using NucleoSpin Gel and PCR clean-up columns. For PCR, VELOCITY high-fidelity DNA polymerase (Bioline) was used with the manufacturer’s reaction mix supplemented with 500 µM betaine (Sigma-Aldrich). For analysis, 2 µL of the reaction were used for DNA gel electrophoresis (0.8% or 2.0% agarose in TAE, Supplementary Figure 5a).

### Recombineering step 1

Circularization of the amplified oligonucleotide pool (0.8 pmol) with the amplified feature-cassette (0.2 pmol) was performed using NEBuilder HiFi DNA Assembly Master Mix (New England Biolabs) in a total reaction volume of 20 µL at 50 °C for 30 min. For analysis by DNA gel electrophoreses, 10 µL of the reaction were used (0.8% agarose in TAE, Supplementary Figure 5b).

### Recombineering step 2

To amplify selectively the circular product from Step 1, rolling circle amplification was used. First, the annealing mixture was set up (total volume: 5 µL in a PCR tube) using 1 µL of the crude or gel-purified circularization reaction, 2 µL exonuclease-resistant random heptamers (500 µM, Thermo Fisher Scientific), 1 µL of annealing buffer (stock: 400 mM Tris-HCl, 50 mM MgCl_2_, pH = 8.0) and 1 µL of water. For annealing, the mixture was heated to 94 °C for 3 min and cooled down in thermocycler at 0.5 °C/sec to 4 °C. Then, 15 µL amplification mixture were added (consisting of 2.0 µL 10 x phi29 reaction buffer, 2.0 µL 100 mM dNTP mix, 0.2 µL 100 x BSA, 10 mg/mL, and 0.6 µL phi29 DNA-polymerase; all from New England Biolabs). Amplification was allowed to proceed for 12–18 h at 30 °C, followed by heat-inactivation of the enzymes at 80 °C for 10 min. For analysis by DNA gel electrophoresis (0.8% agarose in TAE) 0.5 µL of this reaction was used (Supplementary Figure 5c).

### Recombineering step 3

To release the SICs, 20 U of the restriction enzyme *Bst*XI (New England Biolabs) were added directly to the amplification reaction and the mixture was incubated for 3 hours at 37 °C. Typically, such a reaction yielded 10–20 µg of SICs. For DNA gel electrophoresis, 1 µL was used (Supplementary Figure 5d).

### Estimating recombineering fidelity by NGS (Figures 3 and 5)

The oligonucleotide pools were analyzed by NGS after PCR amplification and after recombineering including unique molecular identifiers (UMIs) for de-duplication (Supplementary Figure 7a). For the PCR amplicons, fragments with UMIs were generated using 200 ng starting material (purified by ethanol precipitation) in 2 cycles of PCR with Herculase II Fusion DNA Polymerase (Agilent Technologies) using an equimolar mixture of P023poolseqNN-primers (1 mM final concentration) in a 25 µL reaction. Cycling conditions were based on the manufactures recommendations (62 °C annealing, 30 sec elongation). The reactions were purified with NucleoSpin Gel and PCR clean-up columns using diluted NTI buffer (1:5 in water) to facilitate primer depletion, and the fragments eluted in 20 µL 5 mM Tris-HCl (pH = 8.5) each. To remove residual primers, 7 µL of eluate were treated with 0.5 µL exonuclease I (*E. coli*, New England Biolabs) in 1 x Herculase II reaction buffer (1 h, 37 °C) and heat-inactivated (20 min, 80 °C). The reaction was used without further purification as input for a second PCR (Herculase II Fusion DNA Polymerase, 30 cycles, 72 °C annealing, 30 sec elongation) to introduce indexed Illumina-TruSeq-like adapters (primer Ill-ONP-P7-bi7NN and Ill-ONP-P5-bi5NN). The products were size selected on a 3% NuSieve 3:1 Agarose gel (Lonza), purified using NucleoSpin Gel and PCR clean-up columns, and quantified on a Qubit Fluorometer (dsDNA HS Assay Kit, Thermo Fisher Scientific) and by qPCR (NEBNext Library Quant, New England Biolabs, LightCycler 480, Roche). SIC pools were processed likewise using tRNA-seqNN and mNeon-seqNN as primers to introduce UMIs. All samples were pooled according to the designed complexity and sequenced on a NextSeq 550 system (Illumina) with 300 cycle paired-end chemistry.

### Estimating recombineering fidelity by NGS (Figure 4)

We sequenced the oligonucleotide pool after PCR amplification, and the SIC pool obtained from the recombineering procedure. In the latter instance, fragments compatible with Illumina NGS were generated digesting the products of rolling circle amplification with *Bts*^α^I (55 °C, 90 min, New England Biolabs) and *Sal*I-HF (37 °C, 90 min, New England Biolabs). The fragments were column-purified, diluted to 100 ng/µL and blunted using 1 U/µg mung bean nuclease under the appropriate buffer conditions (New England Biolabs). The DNA fragments of 150–200 bp length were gel-extracted on 3% NuSieve 3:1 Agarose (Lonza). Both samples were sequenced by GATC Biotech AG (Konstanz, Germany) using Illumina MiSeq 150 paired-end NGS technology.

### Transformation of self-integrating cassettes

For transformation of individual self-integrating cassettes (SICs) or SIC pools, Cas12a-family proteins were transiently expressed by making frozen competent cells using either yeasts strains with *GAL1*-controlled Cas12a proteins grown in YP or SC medium containing 2% (w/v) raffinose and 2% (w/v) galactose as carbon source. For transformation^5^, the heat shock was extended to 40 min and no DMSO was added. Recovery of cells that required selection for dominant antibiotic resistance markers (G-418, hygromycin B and clonNAT^6^) was allowed for 5–6 hours at room temperature in YP-Raf/Gal or YPD to proceed prior to plating them on corresponding selection plates.

SIC pools were transformed at a total of 1 µg per 100 µL of frozen competent yeast cells (approximately 2 x 10^8^ cells). Per library approximately 5 of such transformation reactions were combined corresponding to a yeast culture volume of 50 to 100 mL (OD_600_ = 1.0) to generate the competent cells. The number of transformants per library was calculated from serial dilutions. Replica-plating on selective plates was used to exclude transiently transformed clones. After outgrowth, libraries were harvested in 15% glycerol and stored at – 80 °C. For subsequent experiments including genotyping, approximately 10,000 cells per clone were inoculated in YPD, diluted to OD_600_ = 1.0 (approx. 50 mL of culture), and grown over-night. If necessary, a second dilution was performed to obtain cells in exponential growth phase.

For co-integration experiments using individual SICs, 1 µg DNA per SIC and condition was transformed using 50 μL competent yeast cells. Colony number and fluorescence images were acquired after the sample had been spread onto selective plates. Potential co-integrands were tested by replica-plating, streaking and fluorescence microscopy.

Each transformation mixture was split into two parts containing 1/20 (libA) or 19/20 (libB) of the volume. The largest sample was plated onto four 25 x 25 cm square plates with YPD + G-418. No replica-plating was performed before the libraries were cryo-preserved in 2.5 mL, 10 mL, and 50 mL 15% glycerol respectively.

For libraries libC, and the small nuclear library (based on P1), the transformation mixture was plated onto two 25 x 25 cm plates with YPD + hygromcycin B.

### Fluorescence microscopy

Cells were inoculated at an OD_600_ = 0.5 per condition in 5 mL low-fluorescent SC medium (SC-LoFlo^7^) from cryopreservation stocks and grown overnight, followed by dilution to OD_600_ = 0.1 in 20 mL SC-LoFlo the next morning and imaging during mid-exponential growth in the afternoon. Cells were attached to glass-bottom 96-well microscopy plates (MGB096-1-2-LG-L, Matrical) using concanavalin A coating^8^. High-resolution fluorescence micrographs were taken on a Nikon Ti-E epifluorescence microscope equipped with a 60 x ApoTIRF oil-immersed objective (1.49 NA, Nikon), a 2048 x 2048 pixel (6.5 µm), an sCMOS camera (Flash4, Hamamatsu) and an autofocus system (Perfect Focus System, Nikon) with either bright field, 469/35 excitation and 525/50 emission filters or 542/27 excitation and 600/52 emission filters (all from Semrock except 525/50, which was from Chroma). For each condition, a z-stack of 10 planes at 0.5 µm distance was acquired each with a bright field, a short (75% excitation intensity, 10 ms) and a long fluorescence exposure (100% excitation intensity, 100 ms) regimen. For display, the fluorescent image stacks were z-projected for maximum intensity, and cell boundaries taken from out-of-focus bright field images. For imaging cells in Figure 3 (small nuclear pools), cells were inoculated from cryopreservation stocks and grown overnight in selective synthetic media (SC with monosodium glutamate and hygromycin B). On the next morning, the cells were diluted in the same medium and grown to mid-exponential phase. Z-stacks were acquired using 17 planes and 0.3 µm spacing between planes.

### Fluorescence-activated cell sorting (FACS)

A homogenous population of small cells (mostly in G1 phase of the cell cycle) was selected using forward and side scatter. Single cells were sorted according to fluorescence intensity using a FACS Aria III (BD Diagnostics) equipped for the detection of green fluorescent proteins (excitation: 488 nm, long pass: 502LP, bandpass: 530/30). We first isolated cells (3 million in total), which represented roughly the 30% most fluorescent cells in library #1.1 (Supplementary Table 2) as judged by comparison to cells from strain ESM356-1 which was used as negative control. The population of fluorescent cells was then grown to exponential phase and sorted into eight fractions (bins) of 125,000 cells each (except for 62,500 cells sorted into bin 8) using bin sizes of roughly 5% (bin 1), 20%, 20%, 20%, 25%, 5%, 5%, 1% (bin 8) according to the log_10_-transformed intensity of fluorescence emission of small (G1) cells. Sorted pools were grown over-night and the cells were harvested for genomic DNA extraction and target enrichment NGS by Anchor-Seq.

### Library characterization by Anchor-Seq

To determine cassette integration sites in CASTLING libraries, we used a modified Anchor-Seq protocol^9^: Libraries #1.1, #1.2 and #1.3 (Figure 4) were prepared with vectorette bubble adapters (vect_illumina-P5 and vect_illumina-P7) that themselves contained barcodes for multiplexing several samples in the same sequencing run. For all other libraries (Figure 3 and Figure 5), the adapters contained UMIs to account for PCR bias during NGS library preparation (Supplementary Figure 7b); the barcodes for multiplexing were introduced at the stage of the Illumina sequencing adapters. Genomic DNA (gDNA) was isolated from a saturated overnight culture (approximately 2 x 10^8^ cells) using YeaStar Genomic DNA Kit (Zymo Research). Genomic DNA (125 µL at 15 ng µL^−1^ in ultrapure water) was fragmented by sonication to 800–1,000 bp in a microTUBE Snap-Cap AFA Fiber on a Covaris M220 focused ultrasonicator (Covaris Ltd.). In our hands, 51 sec shearing time per tube, a peak incident power of 50 W, a duty factor of 7% and 200 cycles per burst robustly yielded the required size range. Adapters were prepared by combining 50 µM of the respective Watson- and Crick-oligonucleotides (Supplementary Table 6). Each mixture was heated up to 95 °C for 5 min followed by cooling to 23 °C in a large water bath over the course of at least 30 min. Annealed adapters were stored at –20 °C until use. We prepared an equimolar mixture of annealed adapters that contained either none, one, or two additional bases inserted after the UMI (halfY-Rd2-Watson and halfY-Rd2-NN-Crick) to increase heterogeneity of the sequencing library. The fragmented genomic DNA (55.5 µL) were end-repaired and dA-tailed (NEBNext Ultra End Repair/dA-Tailing Module, New England Biolabs) and ligated to 1.5 µL of the 25 µM annealed adapter mix (NEBNext Ultra Ligation Module, New England Biolabs). Products larger than 400 bp were purified by gel excision (using NuSieve, described above) and eluted in 50 µL 5 mM Tris-HCl (pH = 8.5). SIC integration sites were enriched by PCR (NEBNext Ultra Q5 Master Mix, New England Biolabs) using 12 µL of the eluate with suitable pairs of adapter- and SIC-specific primers. Initial denaturation was 98 °C (30 sec), followed by 15 cycles of 98 °C (10 sec), and 68 °C (75 sec). Final extension was carried out at 65 °C (5 min). Reactions were purified using Agencourt AMPure XP beads (0.9 vol, Beckman Coulter). The fragments were further enriched in a second PCR using the custom-designed primers Ill-ONP-P7-bi7NN and Ill-ONP-P5-bi5NN to introduce technical sequences necessary for multiplexed Illumina sequencing. After size-selection by gel extraction (250–600 bp), NGS library concentrations were measured by Qubit Fluorometer (dsDNA HS Assay Kit, Thermo Fisher Scientific) and by qPCR (NEBNext Library Quant, New England Biolabs, LightCycler480, Roche). Furthermore, their size distribution was verified either on a Fragment Analyzer (Advanced Analytical Technologies, Inc) or by gel electrophoresis of the qPCR product. Quantified libraries were sequenced on a NextSeq 500 (for pool C, Deep Sequencing Core Facility) or on a NextSeq 550 sequencing system (both Illumina, 300 cycle paired-end). If necessary, 10–15% phiX gDNA was spiked in to increase sequence complexity.

For MinION nanopore sequencing, the first PCR was carried out as described above for library #1.1 (using 20 cycles) to introduce barcodes for multiplexing FACS bins on the same sequencing run, column-purified, and the NGS library was prepared for 1D sequencing by ligation (SQK-LSK108) according to the manufacturer’s protocols (Oxford Nanopore Technologies). Sequencing was performed on a MinION device using R9.4 chemistry (Oxford Nanopore Technologies). Samples were multiplexed considering the number of different clones present in a pool, bin size, gDNA yield after extraction, and yield of the first PCR.

### Insertion junction sequencing of non-fluorescent cells

Cells from library 1a were grown in selective synthetic media (SC with monosodium glutamate and hygromycin B) for approximately 8 generations, and non-fluorescent cells were sorted into glass-bottom 384-microscopy plates using a FACS Aria III as described under ‘Fluorescence-activated cell sorting (FACS)’. Absence of fluorescence was confirmed by fluorescence microscopy and 60 non-fluorescent clones were pooled and grown overnight to full density. Anchor-Seq amplicons were prepared as described under ‘Library characterization by Anchor-Seq’ using primers NegCells-NNN (Supplementary Table 6). The amplicons were size-selected (∼600 bp) and cloned using the NEB PCR Cloning Kit (New England Biolabs). The resulting amplicons were Sanger-sequenced at Eurofins Genomics (Cologne, Germany).

### Illumina NGS data analysis and read counting

Raw reads (150 bp paired-end) were trimmed and de-multiplexed using a custom script written in Julia v0.6.0 with BioSequences v0.8.0 (https://github.com/BioJulia/BioSequences.jl[https://github.com/BioJulia/BioSequences.jl]).

Read pairs were retained upon detection of basic Anchor-Seq adapter features. Next, these reads were aligned to a reference with all targeted loci using bowtie2^10^ v2.3.3.1. Such references comprised the constant sequence starting from the feature cassette amplified by PCR and 600 bp of the respective proximal genomic sequence of *Saccharomyces cerevisiae* strain S288C (R64-2-1). For off-target analysis, the constant Anchor-Seq adapter features were trimmed off the reads. The remaining variable sequence of the reads was then aligned with bowtie2^10^ to the complete and unmodified genome sequence of *Saccharomyces cerevisiae* strain S288C (R64-2-1). A read pair that aligned to the reference was counted if both reads of the pair were aligned, such that the forward read started at the constant region of the Anchor-Seq adapter-specific primers. In addition, we set the requirement that the inferred insert size was longer than the sequence provided for homologous recombination during the tagging reaction. Counting was implemented using a custom script (Python v3.6.3 with HTSeq 0.9.1^11^ and pysam 0.13). In case UMIs were included in the Anchor-Seq adapter design, they were normalized for sequencing errors using UMI-tools (version 0.5.3)^12^.

For analysis of data obtained from amplicon sequencing (i.e., from PCR and SIC amplification reactions), the reads were either denoised from sequencing errors using dada2 (version 1.5.2)^13^ to evaluate fidelity and abundance or directly aligned with bowtie2 to a reference build from the designed oligonucleotides. Denoised reads were assigned to loci based on the minimal hamming distance to designed oligonucleotides.

### Analysis of nanopore sequencing data and read counting

Nanopore sequencing yields very long reads. Therefore, the reference was assembled as aforementioned but using 2,000 bp of the locus-specific sequences plus the constant sequence of the cassette enriched by the Anhor-Seq reaction. MinION data were base called using the Albacore Sequencing Pipeline Software v2.0.2 (Oxford Nanopore Technologies). For data analysis, a custom script was used to extract and de-multiplex informative sequence segments from all reads based on approximate matching of amplicon features (e.g., the constant region of the vectorette or feature cassette; Julia v0.6.0 with BioSequences v0.8.0, see above). Matching with a Levenshtein distance of 1 was sufficient to discriminate between the barcodes used in this study. Then, the extracted sequence segments were aligned to the reference using minimap2 (v2.2-r409)^14^, using the default parameters (command line option: ‘-ax map-ont’) for mapping of long noisy genomic reads. Only reads that mapped to the beginning of the reference were counted using a custom shell script. The count data for the clones retrieved in each library for cells contained in the individual bins after FACS are provided in Supplementary Table 3.

### Calculation of fluorescence intensity estimates

Fluorescence intensity estimates were calculated as previously described for FACS-based profiling of pooled yeast libraries^15^: Let *b* be a natural number from 1 to 8 indicating one of our 8 FACS bins *B* for which we know the fraction *p_b_* of the total cell population sorted into this bin.

Further, we determined by sequencing for each bin the number of reads *r_g,b_* of an individual genotype *g* (tagged ORF) of all detected genotypes *G*. The obeserved unnormalized cell distribution of *g* is given by:

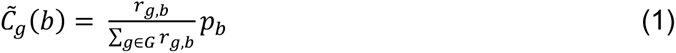

We define the fluorescence intensity estimate for *g* as the empirical mean of *C̃_g_*:

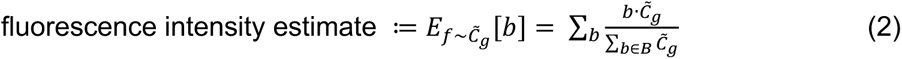

### Calculations and statistical analyses

Statistical analyses were performed using *R* as specified in the scripts or legends.

### Estimation of co-integrand number

We assumed that most co-integrands would result from doubly transformed individuals. So, the number of phenotypic heterozygous individuals (e.g., GFP+ RFP+ or kan^R^ hyg^R^) represents half of the co-integrands if both feature cassettes that were transformed at equimolar ratios have an equal probability of being taken up with the likes of them (i.e., GFP+ GFP+ and RFP+ RFP+) as with each other. Further, we assumed that the fluorescent protein or the antibiotic resistance marker present in the feature cassette had no or only a minor impact on integration efficiency.

### Calculation of copy number changes during RCA

Copy numbers (UMI counts) were normalized to the median UMI frequency in each sequencing experiment and the Gaussian kernel density estimate plotted. Fold changes were calculated as normalized UMI counts after RCA divided by normalized UMI counts after PCR for each oligonucleotide.

### Software and figure generation

Proportional Venn diagrams were generated using eulerAPE^16^. Analyses were performed using R v3.4.1/v3.5.1 with Biostrings v2.44.2^17^ and data.table v1.10.4/v1.11.4. Plots were generated using ggplot2 v2.3.0 and figures were made using Apple Keynote 8.2.

### Data Availability

Raw sequencing data has been deposited on heiDATA (https://dx.doi.org/10.11588/data/L45TRX[https://dx.doi.org/10.11588/data/L45TRX]).

Plasmids and plasmid maps are available upon request. The source data underlying Figures 1b–d, 3b–i, 4a–c, 5b, c, e and Supplementary Figures 1, 2, 3a–c, 4, 5a–d, 8, 9 and 10 are provided as Source Data file.

### Code Availability

The source code of the *R* shiny application for oligonucleotide design is available from our github repository (https://github.com/knoplab/castR/tree/v1.0[https://github.com/knoplab/castR/tree/v1.0]).

## Author contribution

M.K. conceived the project. M.K., B.C.B., K.H. and M.M. designed the experiments and B.C.B., K.H., M.M., and D.K. performed the experiments. E.L. and U.S. contributed methods. K.H., B.C.B., M.K., and M.M. analyzed the data. M.K. and B.C.B. wrote the manuscript. All authors read and approved the final manuscript.

## Acknowledgements

The authors wish to thank Ilia Kats, Cyril Mongis, and Krisztina Gubicza for help with IT infrastructure and experiments. We acknowledge support by the Deutsche Forschungsgemeinschaft (DFG KN498/12-1), the state of Baden-Württemberg through bwHPC for high-performance computing and SDS@hd for data storage (grant INST 35/1314-1 FUGG), and the Dietmar Hopp foundation. K.H. was supported by a HBIGS graduate school fellowship. We also acknowledge help from the Flow Cytometry Core Facility at ZMBH, and the Deep Sequencing Core Facility of the University of Heidelberg, both of which are supported by the CellNetworks cluster of excellence.

## SUPPLEMENTARY INFORMATION

### SUPPLEMENTARY NOTES

#### Supplementary Note 1. Design of an oligonucleotide pool for C-terminal tagging of yeast ORFs

The design principles described below are implemented in *R* (https://github.com/knoplab/ <u>castR/tree/v1.0</u>). For oligonucleotide design the annotated yeast reference genome was retrieved from the *Saccharomyces* Genome Database^1^ (www.yeastgenome.org, November 2016), which contained 5,743 chromosomal yeast ORFs. This process consists of three stages. In stage 1 the CRISPR target sequences are retrieved and evaluated; in stage 2 the corresponding oligonucleotides are constructed *in silico* and evaluated; and in stage 3 the selection of oligonucleotides is further confined to optimally use pool synthesis capacity.

##### Stage 1

For C-terminal tagging, all protospacer-adjacent motifs (PAMs) of the selected CRISPR-Cas12a endonuclease within ±30 nt around the STOP codon were retrieved. For tagging with FnCas12a, these PAMs were TTV and TYN in the first genome-wide pool design and TTV in the second genome-wide pool design. If a PAM was followed by another PAM, only the downstream PAM was considered to obtain non-adjacent CRISPR target sequences (‘spacers’) only. In the next step, the 20 nt long target sequences were extracted. Target sequences that contained runs of more than five thymidine bases were removed to avoid pre-mature termination of crRNA transcription by RNA polymerase III. Spacers prematurely abrogating transcription by ending with T or TT were not removed because 18 nt spacers can still be functional^2^.

For the remaining spacers, we established (1) whether the genomic target sequence would be removed by SIC integration, i.e. whether it spanned the STOP codon, and (2) the risk of off-target cleavage. This risk was assessed as the number of genome-wide occurrences of these sequences (including PAM-free sites) permitting mismatches in the spacer according to the following rules: maximum 1 base mismatch in the crRNA seed, i.e. the first eight PAM-proximal nucleotides, and/or maximum 3 mismatched bases in the entire target sequence. In total, 37,438 TTV sites and 45,445 TYN sites were evaluated during the first genome-wide pool design.

##### Stage 2

Oligonucleotides are constructed as outlined in Supplementary Figure 6. In brief, the length of each homology arm is established first. Any bases potentially removed from the open reading frame (ORF) or the 3’ untranslated region (3’ UTR) after target cleavage by the CRISPR endonuclease are reconstituted by the homology arms after SIC integration.

Next, the oligonucleotides are assembled starting with the last ∼30 nt of the crRNA expression unit, i.e. of the *SNR52* promoter (first genome-wide pool design) or the tG(GC)F2 tRNA site (second genome-wide pool design). In both designs, the next element is the direct repeat sequence of FnCas12a and the target-specific spacer sequence determined in stage 1. Following, the 3’ homology arm, a restriction site used in the recombineering procedure (*Bst*XI in both designs), the 5’ homology arm (without the STOP codon) and finally an in-frame linker, such as a Thr-Ser linker or a S1/S3 linker. The linker serves as generic recombineering site with the feature cassette encoding the desired reporter gene.

After *in silico* design, oligonucleotide sequences that contain more than one of the endonuclease restriction sites are removed. Therefore, it is advised to choose a restriction enzyme with low restriction site frequency at the desired tagging site. For C-terminal tagging in yeast, we considered *Bst*XI or *Ngo*MIV in the first pool design (16,005 remaining TTV candidates; 18,720 remaining TYN candidates) and *Bst*XI in the second pool design (32,602 remaining TTV candidates).

##### Stage 3

Since the number of candidate oligonucleotide designs typically exceeds the synthesis capacity for a single array-based oligonucleotide pool, a selection procedure for pool assembly is required.

In the first genome-wide pool design, this was effected as follows: All ORFs that were targeted by a single oligonucleotide only were preserved regardless of the following criteria. For all other ORFs, oligonucleotides that contained a spacer with low off-target estimate and/or the ability to remove the genomic target by SIC integration were preferred. The final pool contained 12,472 oligonucleotide sequences for tagging 5,664 ORFs.

In the second genome-wide pool design, we tried to consider the parameters previously identified to improve tagging success. Therefore, oligonucleotide candidates were ranked based on a bonus-system. For each ORF, the three oligonucleotides with the highest bonus were selected (17,691 oligonucleotides). We also included designs that had proven repeatedly successful in the first genome-wide pool design, summing up to 18,752 sequences. Another 8,248 candidate oligonucleotides were included by random draw to use up the remaining synthesis capacity of this pool (27,000 oligonucleotides) for tagging 5,940 ORFs.

#### Supplementary Note 2. CASTLING and gene drive

##### Circumstances under which CASTLING can generate a gene drive-system

The co-occurrence of a gene with a molecular machinery that is suited to propagate itself along with that gene from one chromosome to a homologous one, leading to homozygotization of the gene locus with the propagation machinery biases the inheritance of this gene within sexually reproducing populations towards much higher transmission frequencies than expected from sexual inheritance (‘gene drive’). Importantly, the strength of the gene drive depends primarily on the fidelity and efficiency by which the element is copied, and less on the nature of the gene that is driven. In each generation, genes that decrease the fitness of an individual can spread just as efficiently as genes that confer e.g. resistance to antibiotics, leading to potentially harmful or undesired effects on the host population (‘population replacement’^3, 4^, spread of resistances^5^).

Artificial genetic elements that combine the expression of an RNA-programmable endonuclease (i.e. Cas9 or Cas12a) and a gene from which a cognate crRNA is expressed can exhibit gene drive in a process reminiscent of gene conversion^6^.

CASTLING bears the risk of accidentally crating such an artificial drive system if one of the ‘self-integrating’ cassettes is targeted to the neighborhood of a stably integrated, actively expressed Cas12a-endocing gene (Supplementary Figure 11). In such a constellation, the crRNA expressed from the gene engineered by CASTLING could cleave the homologous chromosome and thereby induce repair of the damaged locus with the information of the engineered chromosome. If the CRISPR endonuclease-encoding locus is copied along with the repair template during this process, homozygotization of the drive system is complete. Thus, the likeliness and the strength of the drive depends on the distance between the Cas12a and the crRNA gene, i.e. on the amount of sequence between them that is homologous to the sequence on the homologous chromosome. The exact distance that will eliminate gene drive capabilities is not known, but given the fact that already 36 bp in *S. cerevisiae* (which may be different for other hosts) are sufficient to promote homologous recombination efficiently, we expect that the ability to ‘drive’ will be lost or strongly reduced if the Cas12a gene and the crRNA gene are separated by more than a few hundred base pairs of sequence that is homologous to the other chromosome.

##### Safety and security measures

Based on these considerations we strongly recommend the following safety measures that are effective to prevent the accidental creation of an artificial genetic element that exhibits gene drive using CASTLING in sexually reproducing organisms:

- The Cas12a endonuclease can be contained on an episomal plasmid that is not prone to integrate into the genome. Plasmids that contain counter-selectable markers such as *URA3* can be used to enforce the loss of the Cas12a after library preparation.
- In case a strain with a genomically integrated Cas12a gene is used, the SICs should not target sequences in proximity to the Cas12a locus. In this situation, occasional spontaneous integration of the plasmids into the genome must be considered and care must be taken that none of the SICs can target the plasmid itself. Creating such crRNAs can be avoided by removing (or masking) all sequences that fulfill these criteria from the host reference genome sequence before starting with oligonucleotide pool design.

If these minimal requirements are met and if all the generated genetically modified organisms (GMOs) are contained properly during the experiment, we believe that CASTLING bears no risk of gene drive creation.

### SUPPLEMENTARY FIGURES

**Supplementary Figure 1.**
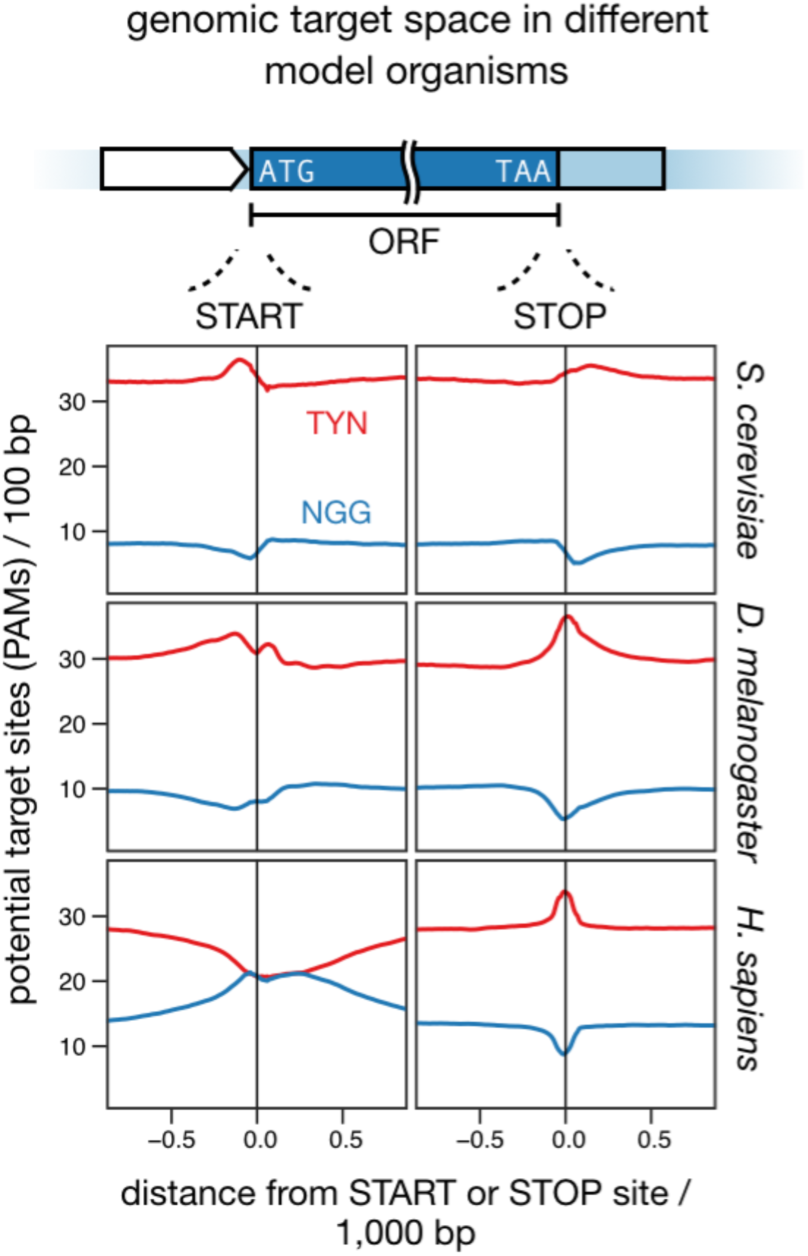
Genomic target space in different organisms considering PAM sites from Cas9 and Cas12a endonucleases. The frequency of the protospacer-adjacent motifs (PAMs) for CRISPR-Cas9 (NGG) and CRISPR-Cas12a (TYN) endonucleases per 100 bp (sliding-window, increment 10 bp) ±1 kb around the START (ATG) and STOP codon (e.g. TAA) of open-reading frames (ORFs) in different organisms is shown. As previously reported^7^, GC-content usually drops or peaks at such sites. Source data are provided as a Source Data file.

**Supplementary Figure 2.**
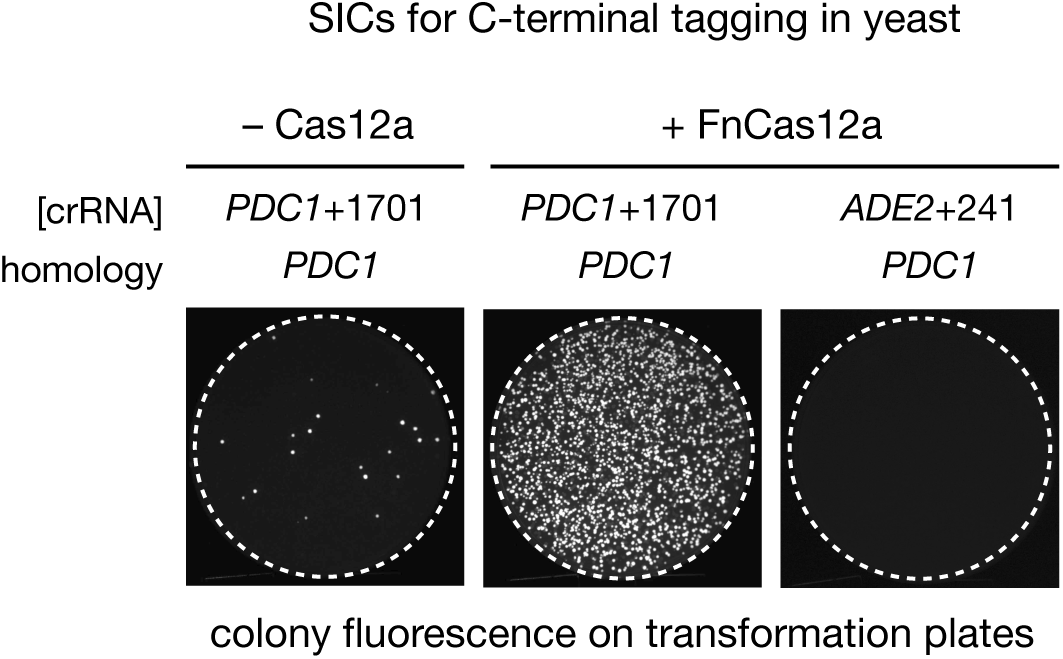
Specific and efficient SIC integration in CASTLING. Transformation of a SIC for C-terminal tagging of *PDC1* using 50 nt homology arms, with and without expression of Cas12a of *Francisella novicida U112* (FnCas12a), or with a crRNA targeting *PDC1* at base +1701, or the *ADE2* locus at base +241 (*ade2* knock-out observed). Source data are provided as a Source Data file.

**Supplementary Figure 3.**
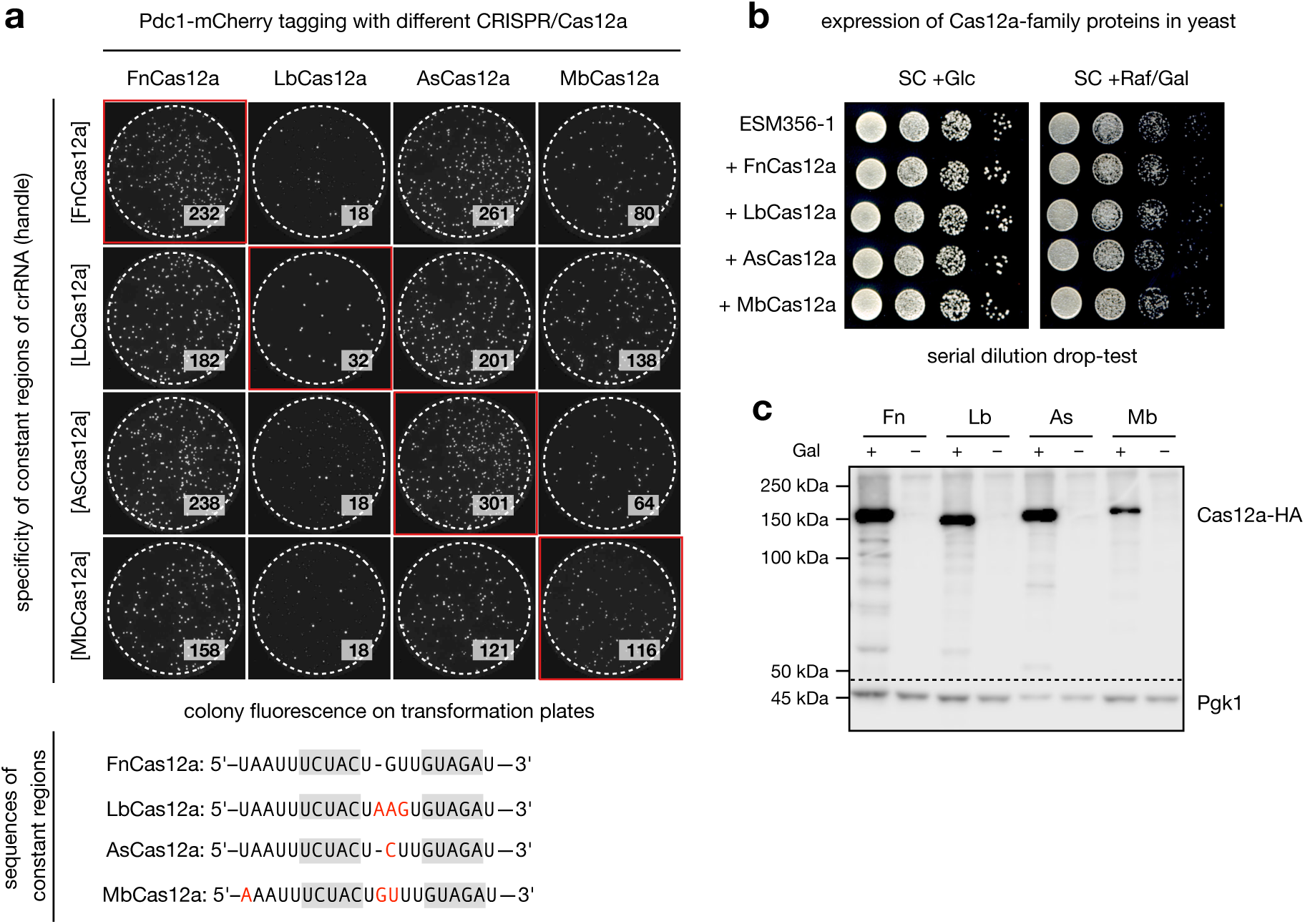
Evaluation of Cas12a-family proteins and expression systems for gene tagging using ‘self-integrating cassettes’ in yeast. (**a**) Combinatorial testing of four different CRISPR-Cas12a-family proteins and four different constant regions (‘handles’) of the crRNA for tagging *PDC1* with a red fluorescent reporter protein (mCherry) by means of [crRNA]*PDC1*+1701 (same PAM, TTTA, in all cases). Colony fluorescence is shown after spread-plating the transformation reaction and outgrowth; numbers indicate the fluorescent colonies growing on the plates after replica-plating. The results demonstrate that integration efficiency of SICs is less dependent on the handle than on the identity of the Cas12a protein. (**b**) Serial dilutions of yeast cultures to test potential toxicity of different CRISPR-Cas12a-family proteins^2^ under expression from a glucose-repressed/galactose-inducible promoter, namely Cas12a of *Francisella novicida U112* (FnCas12a), *Lachnospiriceae bacterium ND2006* (LbCas12a), *Acidaminococcus sp. BV3L6* (AsCas12a), and *Moraxella bovoculi 237* (MbCas12a). None of these Cas12a exhibited toxicity in yeast under these conditions. (**c**) *See page 8*. (**c**) Expression of all tested Cas12a variants was confirmed by immunoblotting (lower panel). Yeast cell extract was normalized to total cell number 3.5 h post induction with 2% galactose. When grown in 2% raffinose as sole carbon-source (−), Cas12a protein levels were below the limit of detection. The Cas12a-family proteins were detected using a hemagglutinin tag and stained by monoclonal antibodies (12CA5, Sigma-Aldrich). Pgk1 (44.7 kDa) was stained as loading control and detected using monoclonal mouse anti-Pgk1 antibodies (R & D Systems, Fisher Scientific). Source data of panels a–c are provided as a Source Data file.

**Supplementary Figure 4.**
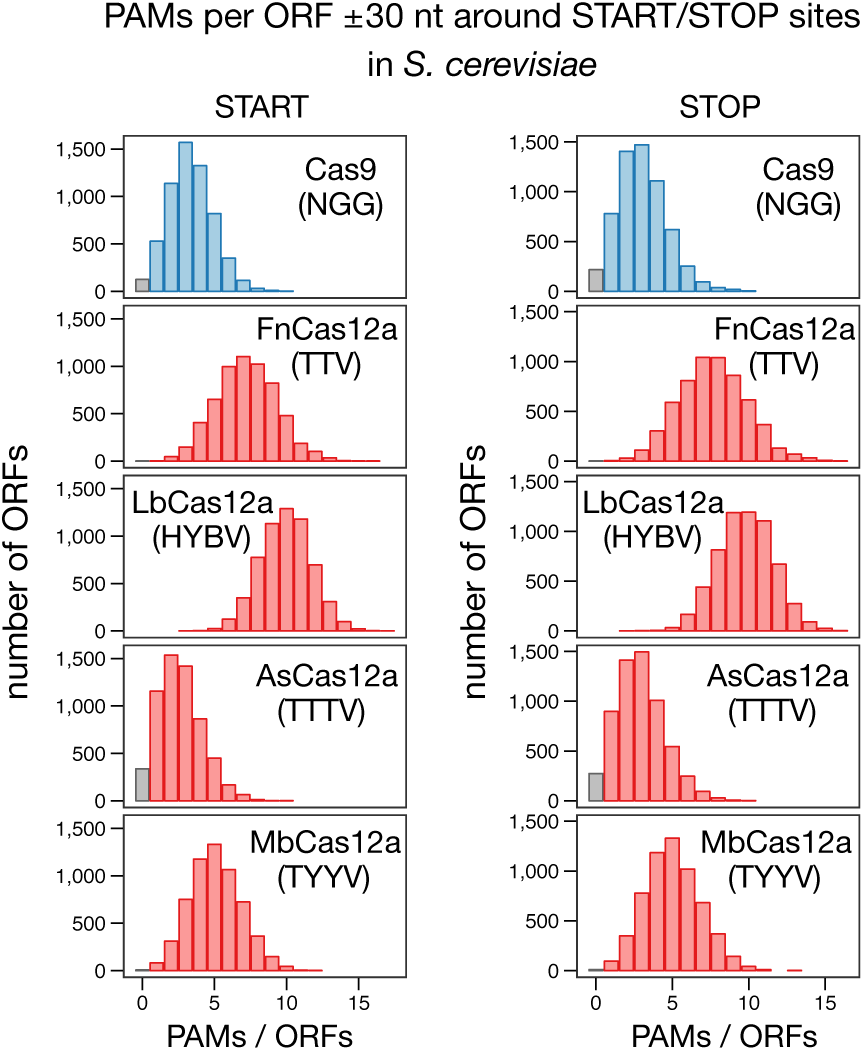
Genomic target space of different CRISPR-Cas12a proteins in yeast. Using CRISPR-Cas12a family proteins, many open-reading frames (ORFs) in yeast can be targeted at multiple sites near (±30 nt) the START or STOP codon. The target space ranges from 83% (AsCas12a) to 99% (FnCas12a) considering all annotated ORFs (retrieved from the *Saccharomyces* Genome Database^1^, www.yeastgenome.org, November 2016). Source data is provided as a Source Data file.

**Supplementary Figure 5.**
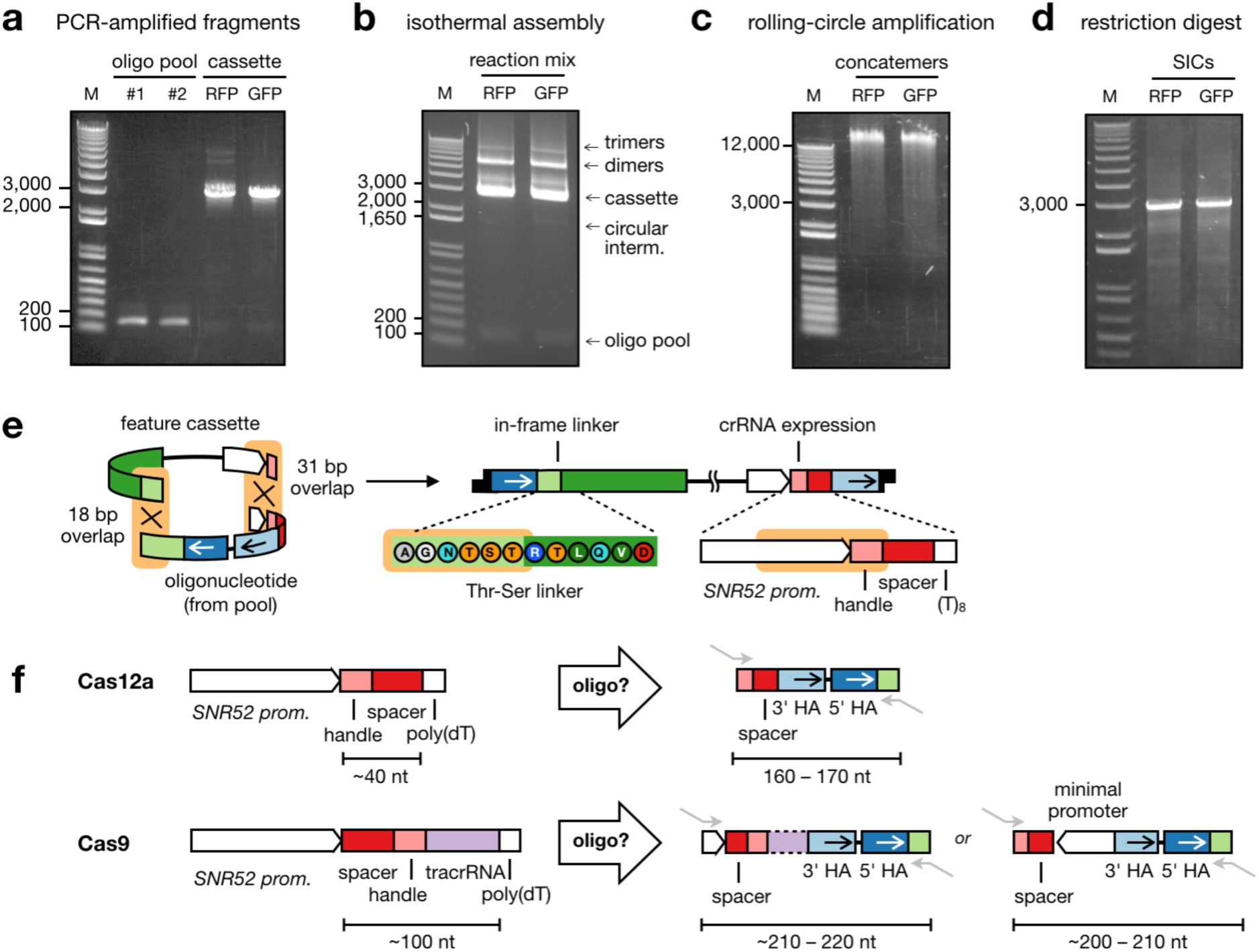
Molecular *in vitro* recombineering procedure to create ‘self-integrating’ cassettes (SICs) from a pool of oligonucleotides. *a–d* displaying agarose gel electrophoresis images of the starting materials, different intermediates and the final SICs. (**a**) The oligonucleotide pool (< 170 bp) and the desired feature cassettes (here: 2–3 kb, but can be shorter or longer) are PCR-amplified (to introduce short homologous overhangs (15–30 bp) for isothermal assembly). (**b**) Molecular recombineering by isothermal assembly yields the ‘self-integrating cassettes’ (SICs) as a covalently-closed circular DNA along with reaction by-products. The circular DNA constitute a minor fraction of the reaction product and for further processing the smallest band is isolated (circular intermediates). (**c**) The circular DNA serves as a template for random-primed isothermal rolling-circle amplification (RCA) to generate very long (>12 kb) linear dsDNA by means of a highly processive DNA-polymerase with strong strand-displacement capabilities such as phi29 DNA-polymerase. Each linear dsDNA is a concatemer repeating a single SIC. (**d–f**) *See page 11*. (**d**) Restriction digestion with a suitable restriction enzyme (here: *Bst*XI) breaks down the long dsDNA molecules into SIC monomers through cleavage of the sequence that linked the homology arms in the designed oligonucleotides. (**e**) Specification of overlap length for isothermal assembly that collapse into an in-frame peptide linker and a functional gene to drive crRNA expression. (**f**) Minimal requirements to provide all locus-specific elements on a single oligonucleotide. For Cas12a, the constant crRNA handle (20 nt) can be part of the primer binding site, and thus the total length of the oligonucleotide ranges 160–170 nt. For Cas9, additional elements must be provided inside the oligo, either a tracrRNA (60 nt) that can be fused with the crRNA to yield a sgRNA (90 nt) or – to reverse the orientation of the elements – a minimal constitutive RNA-polymerase III promoter (35 nt). In total, such oligonucleotides would range 200–220 nt in length. Given current limitations in oligonucleotide synthesis, shorter oligonucleotides (i.e. the Cas12a design) are preferred. Source data of panels a–d are provided as a Source Data file.

**Supplementary Figure 6.**
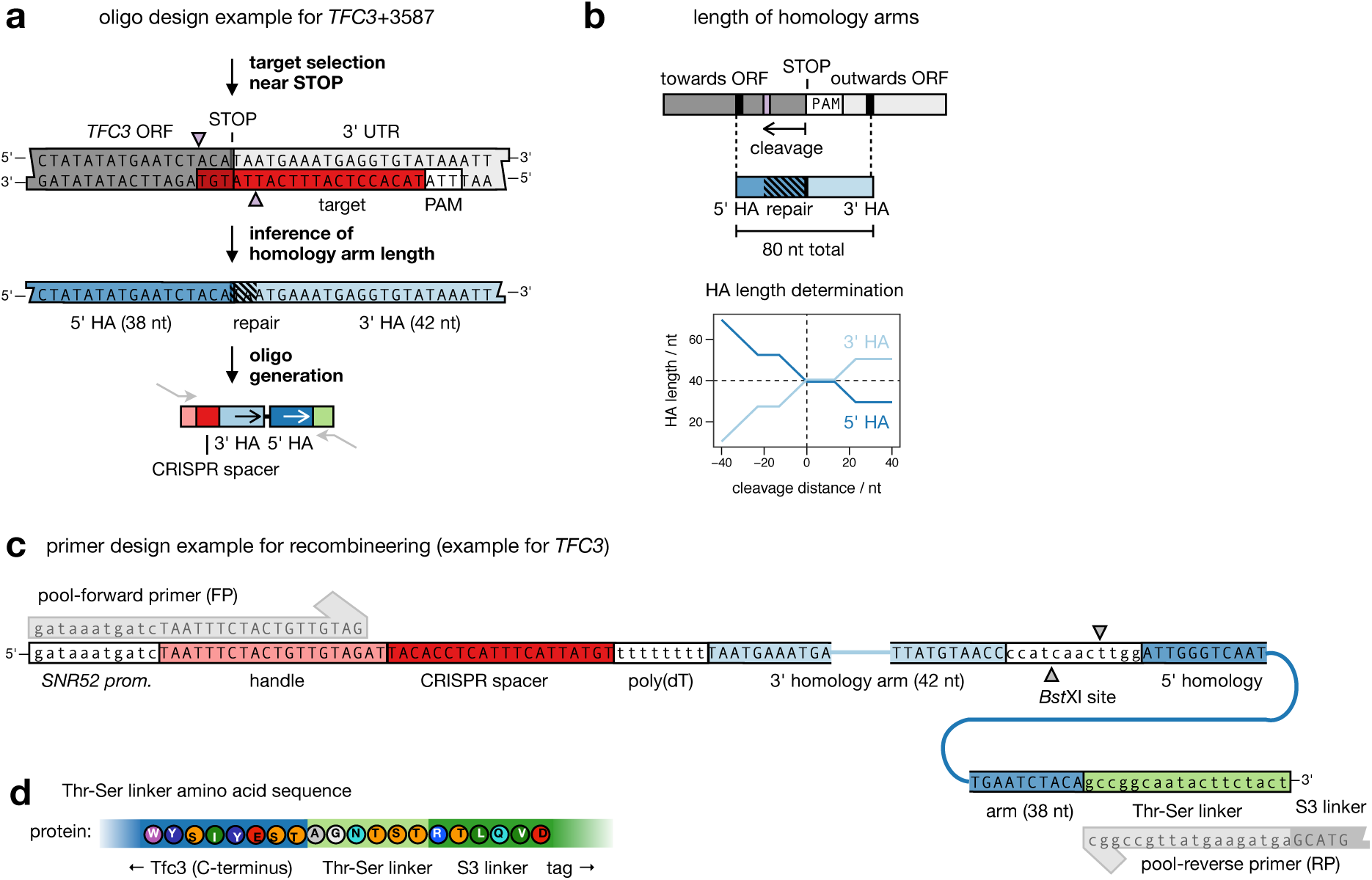
Design example for an oligonucleotide to create a SIC for tagging the gene *TFC3* at position +3587, near the STOP codon. (**a**) Locus-specific elements that are included in the oligonucleotide are the CRISPR spacer complementary to the target sequence for CRISPR-Cas12a, and the homology arms that precisely end and start with the STOP codon for C-terminal tagging. (**b**) The length of homology arms is chosen to always sum up to a total of 80 nt (for space constraints on the oligonucleotide). If the DNA double-strand break occurs upstream of the STOP codon, a part of the 5’ homology arm (HA) serves to reconstitute the lost sequence information and is therefore chosen longer. Experimentally, we determined that homology arms as short as 20 nt still function efficiently to guide correct insertion of the SICs (data not shown). (**c**) Primer design example (as used for the libraries in Figure 4, ID #1.1–1.3 in Supplementary Table 2) for the forward (FP) and reverse primer (RP) of pool amplification. The FP overlaps with the *SNR52* promoter that drives crRNA expression, the RP translates in an in-frame linker between the ORF and the desired tag. (**d**) In-frame peptide linker after C-terminal tagging of Tfc3.

**Supplementary Figure 7.**
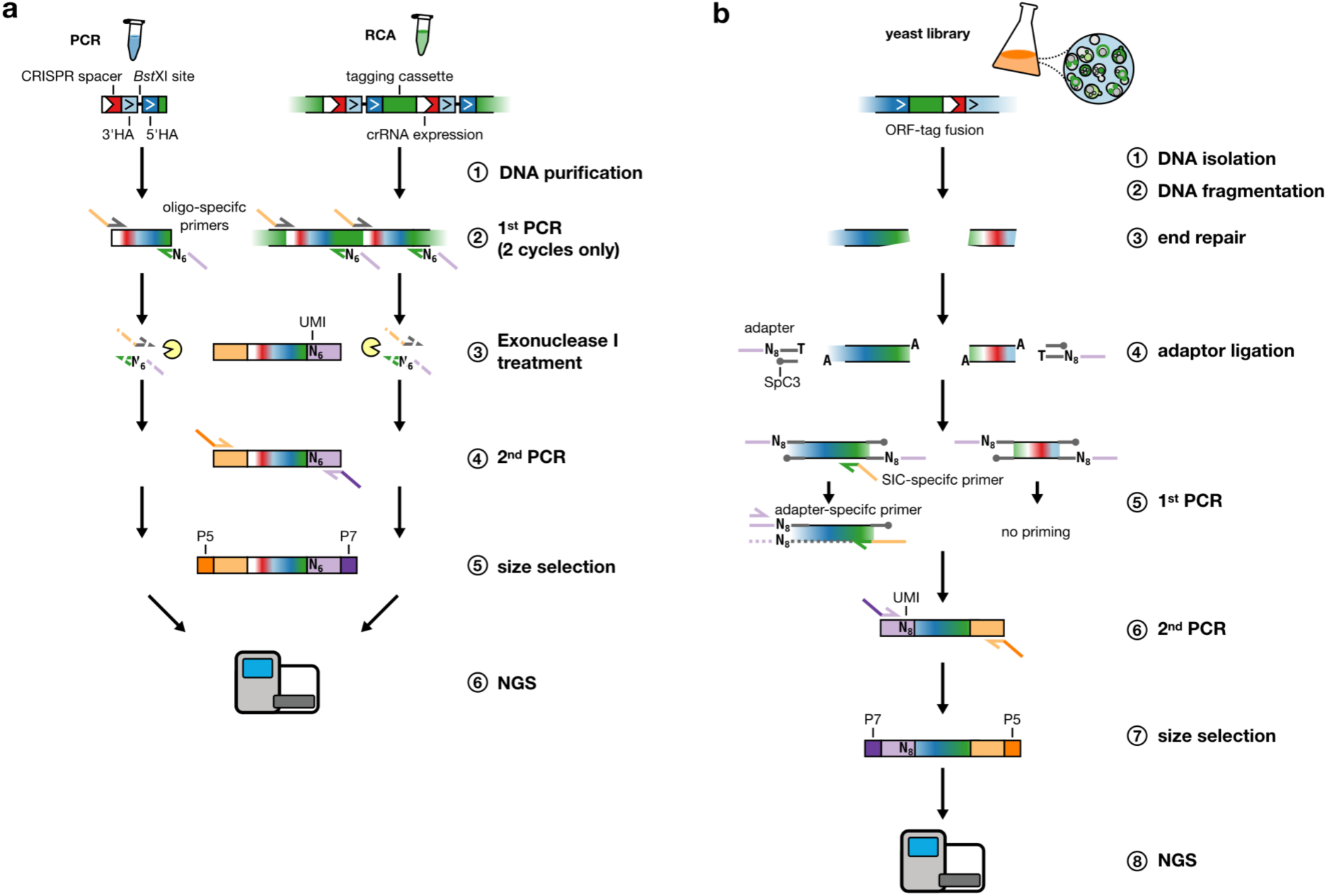
Next-generation sequencing (NGS) library preparation with unique molecular identifiers (UMIs) for molecule counting. (**a**) Quantitative NGS of molecular recombineering intermediates used for CASTLING. The PCR-amplified oligonucleotide pools (left side) or the undigested (concatemeric) SICs yielded by rolling circle amplification (RCA, right side) are quantitated using a random hexameric oligonucleotide sequence (N_6_) as UMI. This N_6_ UMI is incorporated in two rounds of primer annealing and elongation (1^st^ PCR). Unincorporated primers are then destroyed by Exonuclease I treatment, followed by heat inactivation of Exonuclease I. In a final PCR (30 cycles), technical sequences required for Illumina NGS such as P5 and P7 along with barcodes to distinguish different samples are attached. (**b**) For quantitative Anchor-Seq of the yeast libraries, adapters with random octameric nucleotides (N_8_) are ligated to the fragmented and end-repaired genomic DNA. Next, the fragments which contain SIC-derived sequences (the figure shows the junction of the ORF and the tag) are selectively amplified in a PCR with a SIC-specific and a reverse primer. During the initial cycle, the SIC-specific primer generates the binding site of the reverse primer by replication of an asymmetric sequence. This allows for these genomic fragments to be exponentially amplified. An elongation-inhibiting group (Spacer C3, SpC3) on the complementary strand of the ligated adapters prohibits non-specific amplification of fragments without the SIC-specific sequence of interest.

**Supplementary Figure 8.**
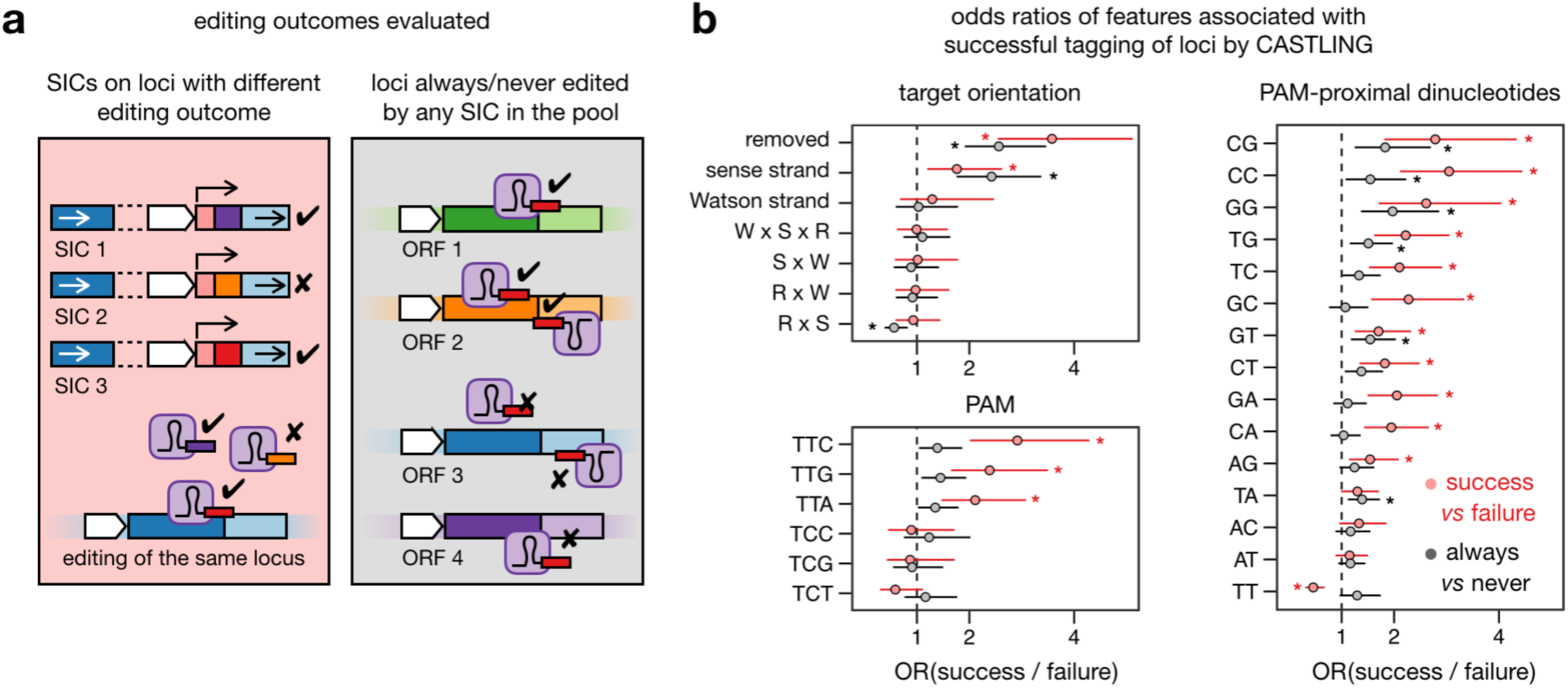
Features of the crRNA and target locus associated with successful tagging. (**a**) Details to the analysis shown in Figure 4d. Two scenarios were considered as represented by red and grey panels. Scenario 1 (red panel) considers all SICs demonstrably present in the SIC pool for the same gene to be tagged, comparing them based on the editing outcome (success: *N* = 2,053 SICs; failure: *N* = 2,488 SICs). Only genes were included in this analysis, that had both, successful and non-successful SICs. Scenario 2 (gray panel) compares tagging success for genes, in which any of the designed SICs promoted (total *N* = 2,885 SICs) or failed (*N* = 3,209 SICs) to tag the gene. (**b**) Odds ratio (OR) based on samples drawn akin to (*a*) based on multiple logistic regression analysis. Asterisks indicate properties for which the impact on odds is significantly (Fisher’s exact test, p < 0.01) different from 1 (no effect); error bars represent 95% confidence intervals). All scenarios displayed were checked for inter-dependencies (indicated as ‘A x B’) also in-between the panels with no effect on the general outcome, but were split for the sake of clarity. ‘Sense strand’ refers to the genomic target occurring on the non-transcribed strand, whereas ‘Watson strand’ is any of the two strands as specified in the yeast gene nomenclature. ‘Nucleosome occupancy at the PAM’ or across the entire ‘target sequence’^8^ and ‘gene expression in YP-Raf/Gal’ (medium to transiently express FnCas12a) or ‘YPD’^9^ (to suppress FnCas12a expression) were also analyzed with no significant impact on tagging success (data not shown). Source data are provided as a Source Data file.

**Supplementary Figure 9.**
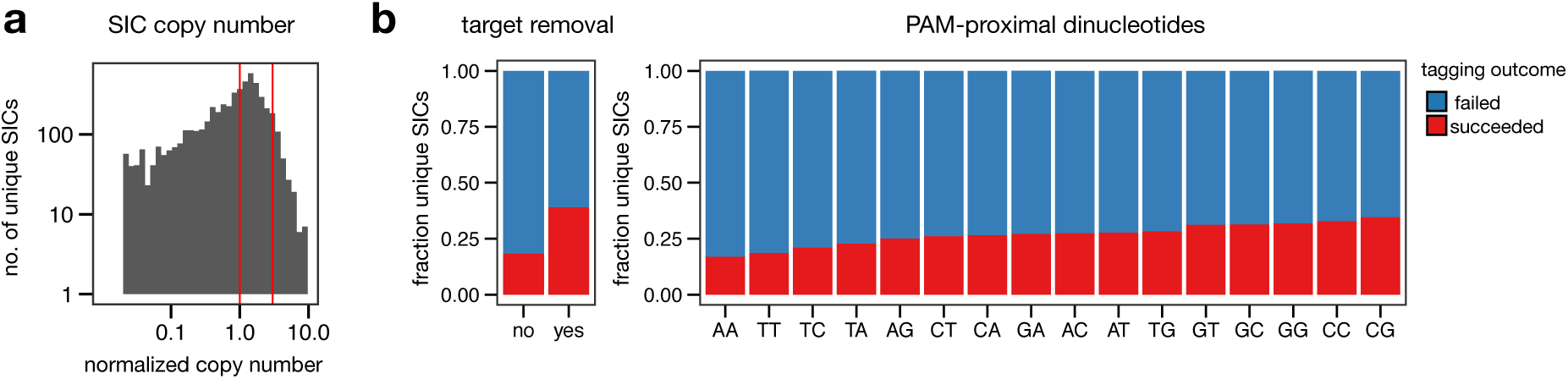
Verification of features of the crRNA and the target locus that are associated with successful tagging. (**a**) Copy number distribution of the SICs in the SIC pool after RCA. Integration success of a particular SIC is likely to be influenced by its copy number in the SIC pool. (**b**) To validate the impact of different features on the tagging success, we restricted the analysis of the impact of different features on tagging success to SICs with similar representation in the SIC pool (1- to 3-fold of the median copy number, region marked with red lines in (a)). Target removal (left panel): Comparison of SICs on the basis whether their integration does remove the target site or not. Influence of PAM-proximal dinucleotides: Comparison of the influence of the PAM-proximal dinucleotides in the crRNA sequence on tagging success (right panel). Both plots demonstrate that tagging success was higher when the rule set (Supplementary Figure 8) was obeyed. Source data are provided as a Source Data file.

**Supplementary Figure 10.**
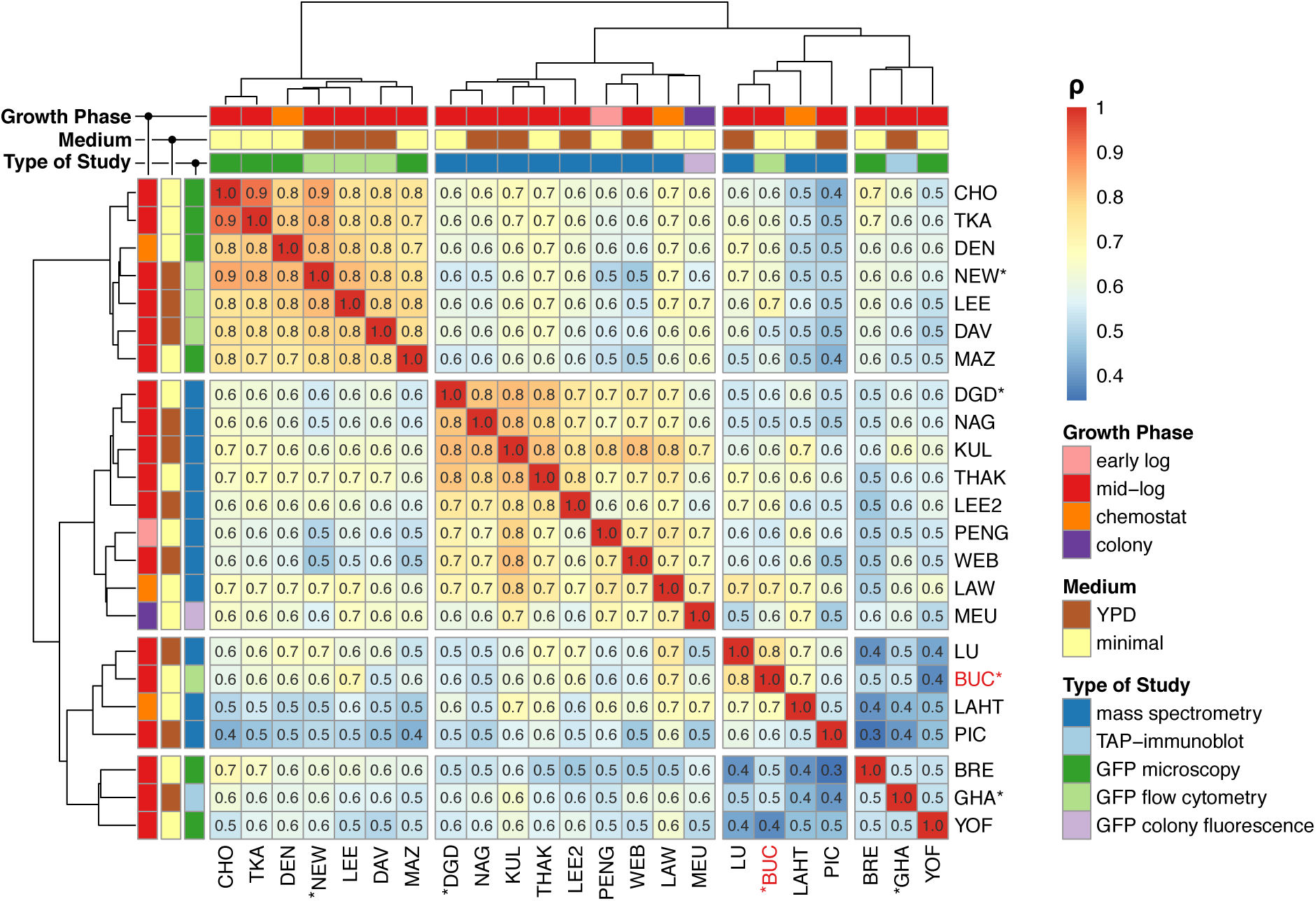
Spearman correlation coefficients (ρ) between protein abundance estimated after phenotyping by FACS in this study (BUC, marked red, Figure 5d–f) and other studies. Study abbreviations as in Ref^10^ except for (MEU)^11^ and this study (BUC). An asterisk indicates studies that appear in Figure 5e. Additional information about each study (growth phase, medium, type of study) were compiled from information in Ref^10^.

Source data are provided as a Source Data file.

**Supplementary Figure 11.**
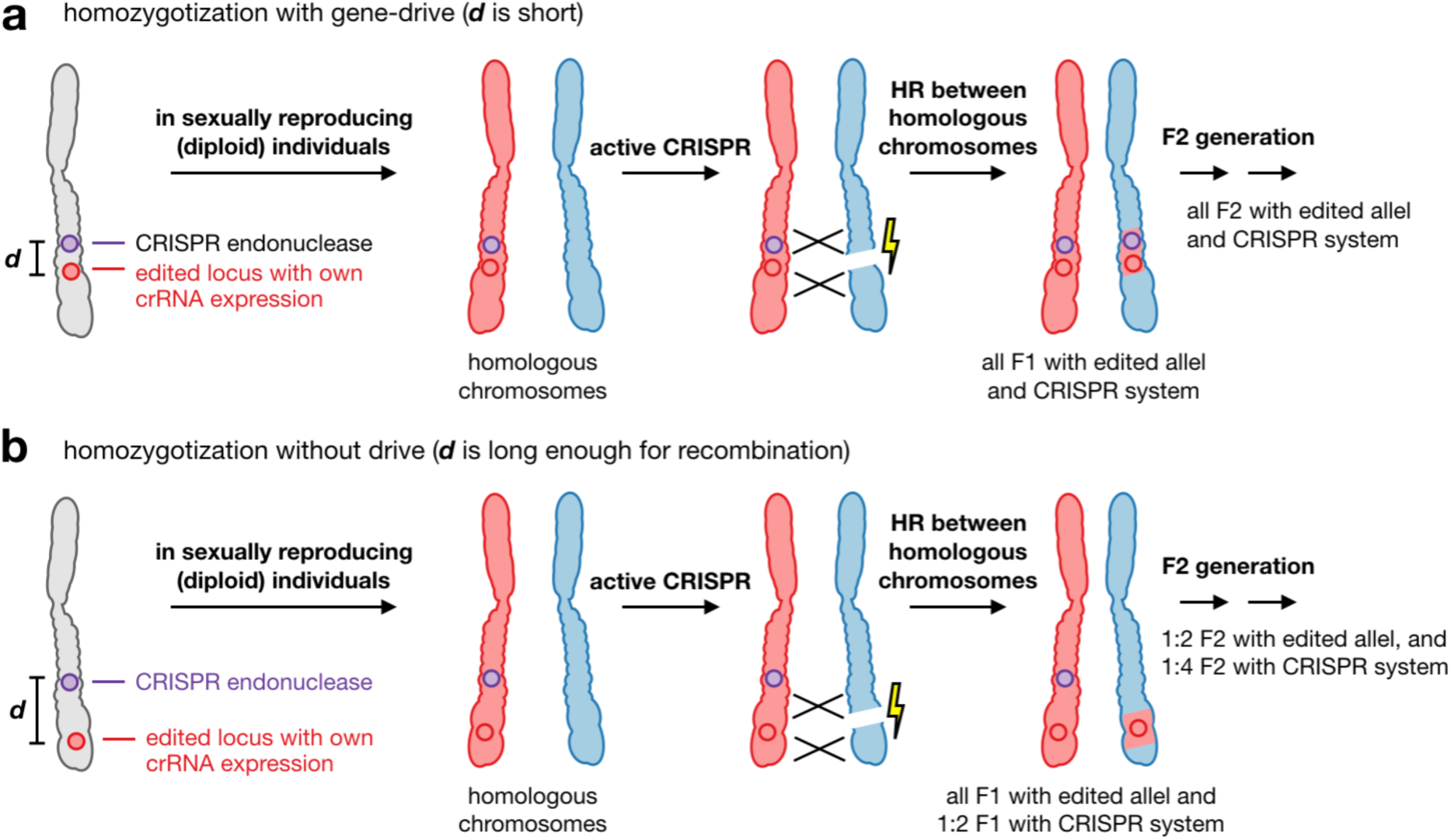
Mechanism of allele homozygotization and potential ‘gene drive’ with stably integrated CRISPR endonucleases (Supplementary Note 1). (**a**) If the distance (***d***) of the crRNA expression site and the endonuclease locus is very short, homozygotization of both loci is highly likely (gene drive). (**b**) With increasing ***d***, the likelihood that both genes are copied to the homologous chromosome decreases.

### SUPPLEMENTARY TABLES

**Supplementary Table 1.** Library statistics for libraries created from pools A, B1 and B2 (Figure 3).

| ID | oligo-nucleotide pool | PCR pool (after amplification) |  |  | feature cassette | SIC pool (after recombineering/RCA) |  |  | clone collections (libraries) |  |  |  |
| --- | --- | --- | --- | --- | --- | --- | --- | --- | --- | --- | --- | --- |
|  |  | primer | reads mapped | identified unique sequences, ORFs (UMI >2) |  | pooled RCA reactions | reads mapped | identified unique sequences, ORFs (UMI > 2) | clones | library stock | reads mapped | identified unique sequences, ORFs (UMI >2) |
| 1a | pool A, 40.0 ng (0.8 pmol) | pool-FP3 + pool-RP3 | 152,011 | 1,200 seq. (76%),<br>206 ORFs (96%) | pKH51 | 2 | 125,015 | 493 seq. (31%),<br>147 ORFs (68%) | 58,000 | YKH231 | 277,026 | 280 seq. (18%),<br>125 ORFs (58%) |
| 1b |  |  | 195,535 | 1,281 seq. (81%),<br>209 ORFs (97%) | pKH51 | 2 | 70,984 | 324 seq. (20%),<br>116 ORFs (54%) | 64,000 | YKH232 | 266,519 | 202 seq. (13%),<br>107 ORFs (50%) |
| 2a | pool B1, 150. pg (3.0 fmol) | pool-FP2 + pool-RP2 | 265,556 | 1,308 seq. (83%),<br>209 ORFs (97%) | pKH51 | 3 | 177,648 | 872 seq. (55%),<br>191 ORFs (89%) | 82,000 | YKH227 | 443,204 | 496 seq. (31%),<br>172 ORFs (80%) |
| 2b |  |  | 253,732 | 1,333 seq. (84%),<br>209 ORFs (97%) | pKH51 | 2 | 225,114 | 601 seq. (38%),<br>179 ORFs (83%) | 81,500 | YKH228 | 478,846 | 309 seq. (19%),<br>144 ORFs (67%) |
| 3a | pool B2, 150. pg (3.0 fmol) | pool-FP3 + pool-RP3 | 148,529 | 1,454 seq. (92%),<br>211 ORFs (98%) | pKH51 | 2 | 44,005 | 825 seq. (52%),<br>177 ORFs (82%) | 58,500 | YKH229 | 192,534 | 233 seq. (15%),<br>127 ORFs (59%) |
| 3b |  |  | 143,308 | 1,435 seq. (91%),<br>211 ORFs (98%) | pKH51 | 2 | 133,050 | 1,082 seq. (67%),<br>196 ORFs (91%) | 32,000 | YKH230 | 276,358 | 265 seq. (17%),<br>135 ORFs (63%) |
| 4a | pool B2, 1.0 ng (20 fmol) | pool-FP3 + pool-RP3 | 179,709 | 1,399 seq. (89%),<br>211 ORFs (98%) | pKH51 | 3 | 60,323 | 1,033 seq. (66%),<br>191 ORFs (89%) | 90,500 | YKH233 | 240,032 | 812 seq. (51%),<br>197 ORFs (92%) |
| 4b |  |  | 202,657 | 1,347 seq. (85%),<br>211 ORFs (98%) | pKH51 | 3 | 52,332 | 964 seq. (61%),<br>189 ORFs (98%) | 95,000 | YKH234 | 254,989 | 835 seq. (53%),<br>197 ORFs (92%) |

**Supplementary Table 2.** Library statistics pool C (Figure 4) and pool D (Figure 5). * = DADA2 de-noising, considering only perfectly matched reads to the reference.

| ID | oligo-nucleotide pool | PCR |  |  | feature cassette | SIC pool (after recombineering, RCA) |  |  | clone collections (libraries) |  |  |  |
| --- | --- | --- | --- | --- | --- | --- | --- | --- | --- | --- | --- | --- |
|  |  | primer | reads mapped | identified unique sequences, ORFs |  | pooled RCA reactions | reads mapped | identified, unique sequences, ORFs | clones | library stock | reads mapped | identified unique sequences, ORFs |
| #1.1 | pool C, 1 x 80 pg (1.5 fmol) | pool-FP1 + pool-RP1 | 1,610,291 | 3,802 seq.* (30%),<br>2,985 ORFs (53%) | pMaM505 (kanMX) | 4 | 3,066,702 | 3,196 SICs* (26%),<br>2,522 ORFs (45%) | 109,000 | YDK553 + YDK557 | 28,574,949 | 2,052 ORFs (36%) |
| #1.2 |  | pool-FP1 + pool-RP1 |  |  | pBeB019 (hphNT1) | 4 | n. d. | n. d. | 100,000 | YDK556 + YDK483 | 28,619,645 | 2,317 ORFs (35%) |
| #1.3 | pool C, 10 x 80 pg (1.5 fmol) | pool-FP1 + pool-RP1 | 18,009,833 | 8,110 seq.* (65%),<br>5,025 ORFs (89%) | pBeB019 (hphNT1),<br>1/10 of PCR | 4 | n. d. | n. d. | 76,500 | YDK559 | 22,476,787 | 2,033 ORFs (36%) |
| LibA | pool D, 1 x 2 ng (40 fmol) | pool-FP2 + pool-RP2 | n.d. | n.d. | pBeB064 (kanMX) | 30 | n.d. | n.d. | 704,000 | YKG7+<br>YKG10+<br>YKG13 | 52,992,365 | 3,786 ORFs (64%) |
| LibB |  |  |  |  |  |  |  |  | 44,000 | YKG5+<br>YKG8+<br>YKG11 | 4,006,055 | 2,869 ORFs (48%) |
| LibC |  |  |  |  | pKH51 (hphNT1) | 2 | n.d. | n.d. | 116,000 | YKH225 | 10,006,829 | 3,011 ORFs (51%) |

**Supplementary Table 3.** Newly characterized ORFs in terms of protein abundance in liquid minimal medium. The detection probability is a function of the relative abundance of a clone in a sorted pool of cells (or ‘bin’) and it is influenced by the selectivity of the phenotype enrichment protocol and the accuracy of genotype identification. For each ORF identified after MinION nanopore sequencing, the read count (> 1 count) and enrichment or depletion (column ‘fold’) in relation to the representation of each ORF in the starting pool, which was characterized using Illumina NGS. Only enriched loci were considered as hits. ORFs for which abundance information was available outside genome-scale experiments^12^ are asterisked.

| bin | enriched loci |  |  | depleted loci |  |  |
| --- | --- | --- | --- | --- | --- | --- |
|  | systematic name | counts | fold | systematic name | counts | fold |
| bin 2<br>(5–24%) |  |  |  | YBR298C-A | 5 | 0.39x |
|  |  |  |  | YJL136W-A | 3 | 0.71x |
|  |  |  |  | YMR175W-A* | 7 | 0.35x |
|  |  |  |  | YOL164W* | 3 | 0.23x |
| bin 3<br>(25–44%) | YGR146C-A | 412 | 3.9x | YGR240C-A | 8 | 0.22x |
|  | YGR204C-A | 61 | 2.0x | YJR151W-A | 146 | 0.24x |
|  | YOL156W* | 18 | 4.4x |  |  |  |
|  | YOR072W-B | 2 | 1.3x |  |  |  |
|  | YPL038W-A* | 17 | 2.7x |  |  |  |
| bin 4<br>(45–64%) | YDR194W-A* | 4 | 4.5x |  |  |  |
| bin 5<br>(65–89 %) | YGL006W-A | 5 | 10.3x |  |  |  |
|  | YML054C-A | 316 | 4.8x |  |  |  |
|  | YMR230W-A* | 8 | 3.1x |  |  |  |
| bin 7<br>(95–99 %) | YBR196C-A* | 570 | 4.2x |  |  |  |

**Supplementary Table 4.** Strains used in this study.

| strain | background | description | purpose | source |
| --- | --- | --- | --- | --- |
| **ESM356-1** | S228C | MATa *ura3-52 leu2Δ1 his3Δ200 trp1Δ63* | auxotrophic laboratory yeast strain | Ref.<sup>13</sup> |
| **YBeB500** | ESM356-1 | *ura3-52::GAL1pr-FnCas12a::NLS::3xHA-CYC1t (natNT2)* | inducible expression in yeast (use discouraged for risk of gene-drive) | this work |
| **YBeB600** | ESM356-1 | *ura3-52::GAL1pr-LbCas12a::NLS::3xHA-CYC1t (natNT2)* | inducible expression in yeast (use discouraged for risk of gene-drive) | this work |
| **YBeB700** | ESM356-1 | *ura3-52::GAL1pr-AsCas12a::NLS::3xHA-CYC1t (natNT2)* | inducible expression in yeast (use discouraged for risk of gene-drive) | this work |
| **YBeB800** | ESM356-1 | *ura3-52::GAL1pr-MbCas12a::NLS::3xHA-CYC1t (natNT2)* | inducible expression in yeast (use discouraged for risk of gene-drive) | this work |

**Supplementary Table 5.** Plasmids used in this study.

| plasmid | backbone | description | purpose | source |
| --- | --- | --- | --- | --- |
| **pY004** | pcDNA 3.1(+) | CMVpr-hFnCas12a::NLS ::3xHA::bGH poly(A) | mammalian expression | Ref.<sup>2</sup> |
| **pY010** | pcDNA 3.1(+) | CMVpr-hAsCas12a::NLS ::3xHA::bGH poly(A) | mammalian expression | Ref.<sup>2</sup> |
| **pY014** | pcDNA 3.1(+) | CMVpr-hMbCas12a::NLS ::3xHA::bGH poly(A) | mammalian expression | Ref.<sup>2</sup> |
| **pY016** | pcDNA 3.1(+) | CMVpr-hLbCas12a::NLS ::3xHA::bGH poly(A) | mammalian expression | Ref.<sup>2</sup> |
| **pRS306N** |  | shuttle vector and yeast integrating plasmid at *ura3-52* conferring nourseothricin resistance |  | Ref.<sup>14</sup> |
| **pMaM486** | pRS306N | GAL1pr-hFnCas12a::NLS ::3xHA-CYC1t | stable genomic integration in yeast, inducible expression | this work |
| **pMaM487** | pRS306N | GAL1pr-hAsCas12a::NLS ::3xHA-CYC1t | stable genomic integration in yeast, inducible expression | this work |
| **pMaM488** | pRS306N | GAL1pr-hMbCas12a::NLS ::3xHA-CYC1t | stable genomic integration in yeast, inducible expression | this work |
| **pMaM489** | pRS306N | GAL1pr-hLbCas12a::NLS ::3xHA-CYC1t | stable genomic integration in yeast, inducible expression | this work |
| **pRS416** |  | shuttle vector and yeast centromeric plasmid conferring uracil prototrophy |  | Ref.<sup>15</sup> |

| (Supplementary Table 5 continued) |  |  |  |  |
| --- | --- | --- | --- | --- |
| plasmid | backbone | description | purpose | source |
| pRS415 |  | shuttle vector and yeast centromeric plasmid conferring leucine prototrophy |  | Ref.<sup>15</sup> |
| pBeB503 | pRS416 | TEF1pr-hFnCas12a::NLS::3xHA-CYC1t | constitutive expression in yeast | this work |
| pBeB603 | pRS416 | TEF1pr-hLbCas12a::NLS::3xHA-CYC1t | constitutive expression in yeast | this work |
| pBeB703 | pRS416 | TEF1pr-hAsCas12a::NLS::3xHA-CYC1t | constitutive expression in yeast | this work |
| pBeB803 | pRS416 | TEF1pr-hMbCas12a::NLS::3xHA-CYC1t | constitutive expression in yeast | this work |
| pBeB501 | pRS415 | GAL1pr-hFnCas12a::NLS::3xHA-CYC1t | inducible expression in yeast | this work |
| pBeB601 | pRS415 | GAL1pr-hLbCas12a::NLS::3xHA-CYC1t | inducible expression in yeast | this work |
| pBeB701 | pRS415 | GAL1pr-hAsCas12a::NLS::3xHA-CYC1t | inducible expression in yeast | this work |
| pBeB801 | pRS415 | GAL1pr-hMbCas12a::NLS::3xHA-CYC1t | inducible expression in yeast | this work |
| pFA6a |  | E. coli plasmid with ampicillin resistance gene ampR |  | Ref.<sup>16</sup> |
| pBeB019 | pFA6a | S3-mScarlet-i-ADH1t (hphNT1) SNR52pr-FnCas12a handle | feature cassette template with a GFP conferring hygromycin resistance | this work |
| pBeB013 | pFA6a | S3-mNeonGreen-ADH1t (hphNT1) SNR52pr-FnCas12a handle | feature cassette template with a GFP conferring hygromycin resistance | this work |
| pMaM505 | pFA6a | S3-mNeonGreen-ADH1t (kanMX4) SNR52pr-FnCas12a handle | feature cassette template with a GFP conferring G-418 resistance | this work |
| pBeB064 | pFA6a | mNeonGreen-ADH1t(kanMX4) SNR52pr-t(GCC)F1-FnCas12a handle | feature cassette template with a GFP conferring G-418 resistance | this work |
| pKH51 | pFA6a | mNeonGreen-ADH1t(hphNT1) SNR52pr-t(GCC)F1-FnCas12a handle | feature cassette template with a GFP conferring hygromycin resistance | this work |

**Supplementary Table 6.**
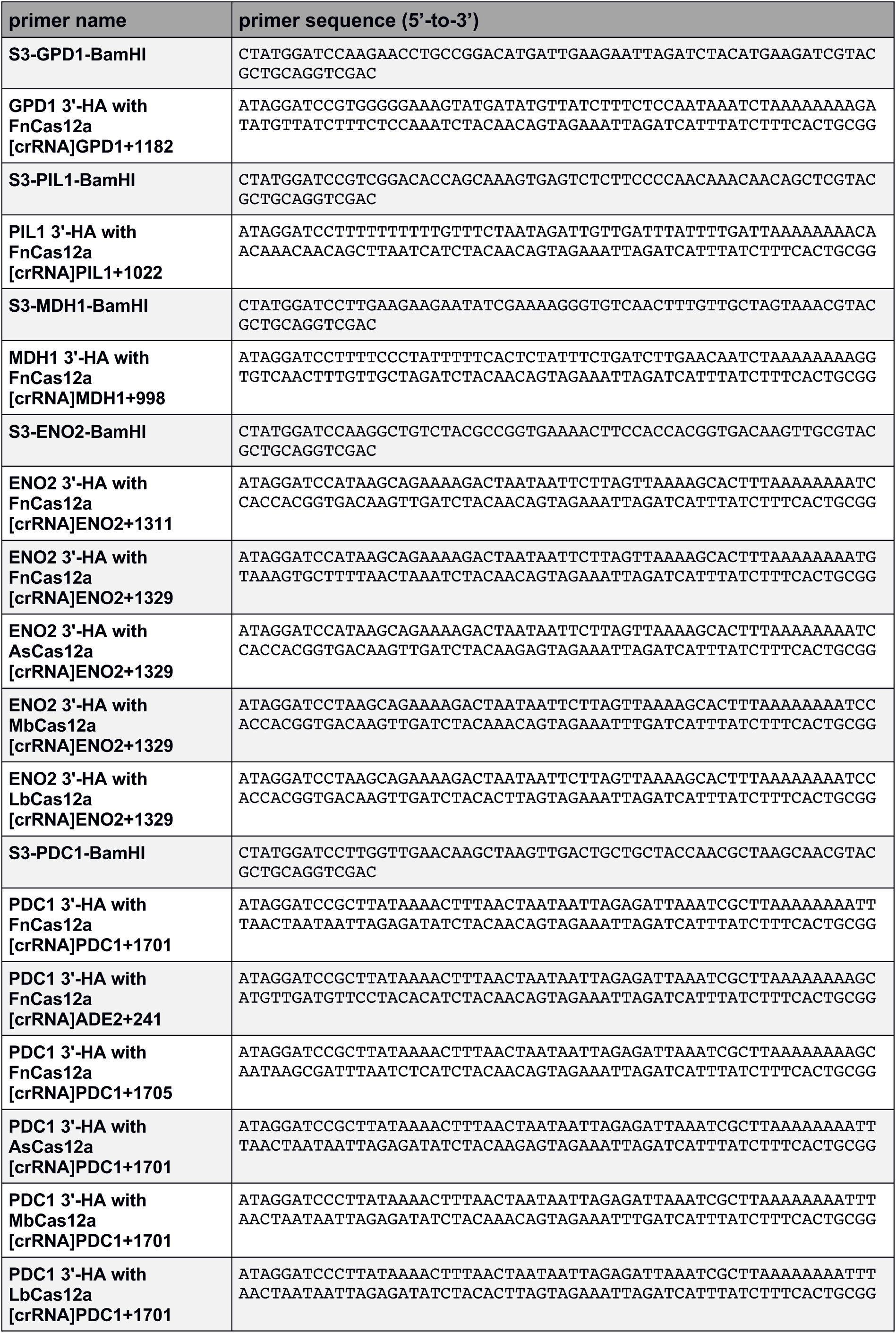
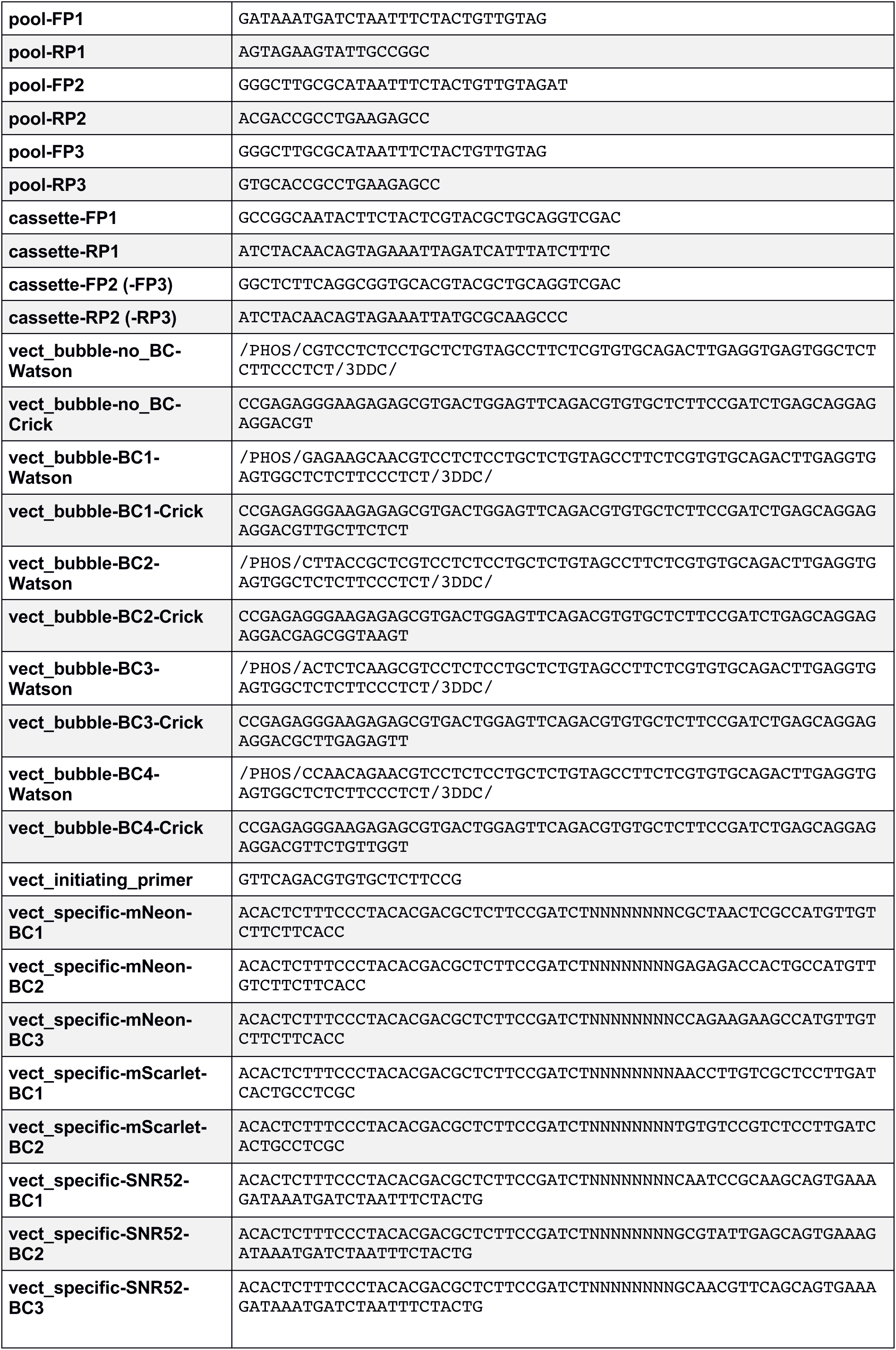
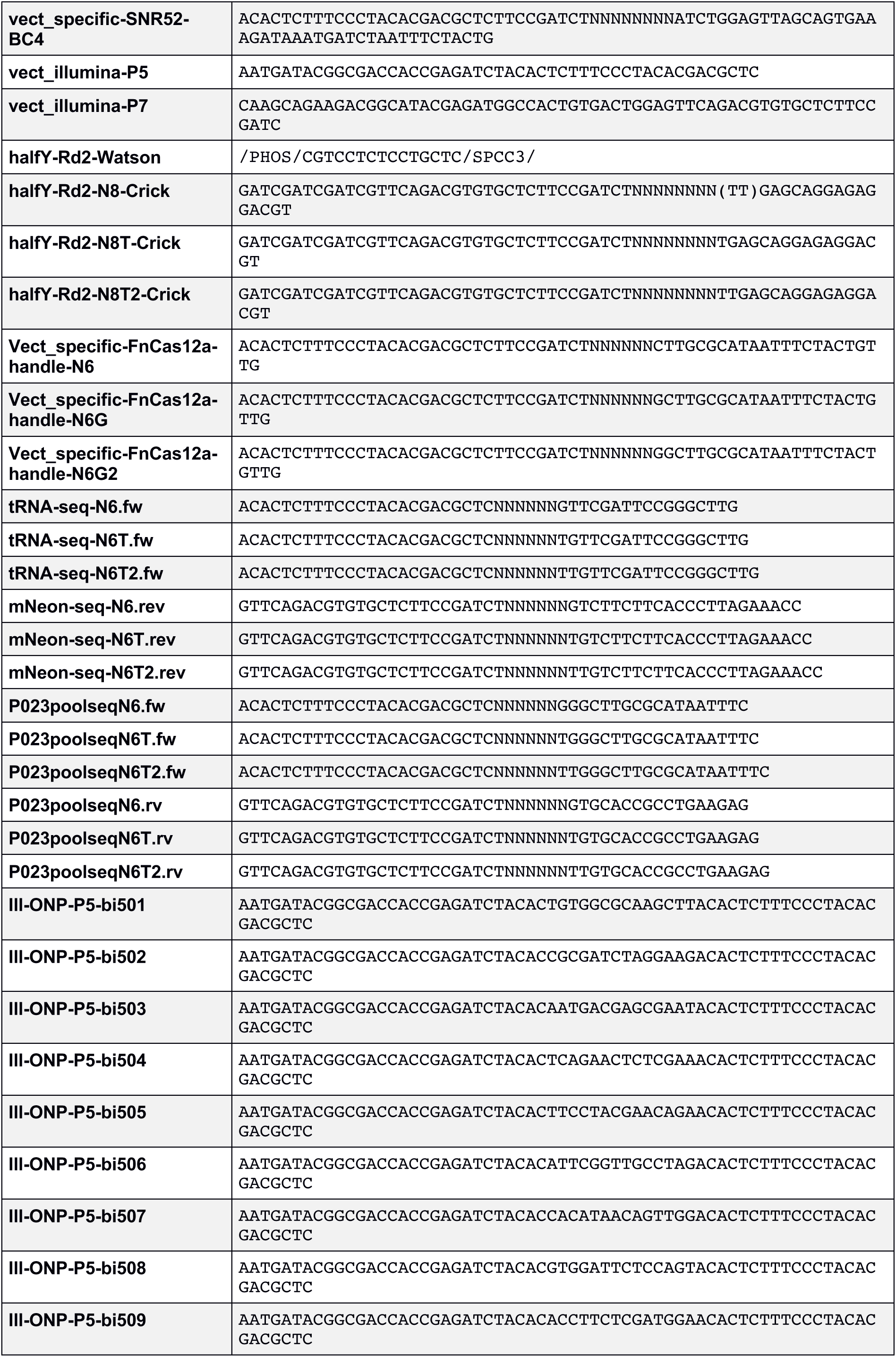
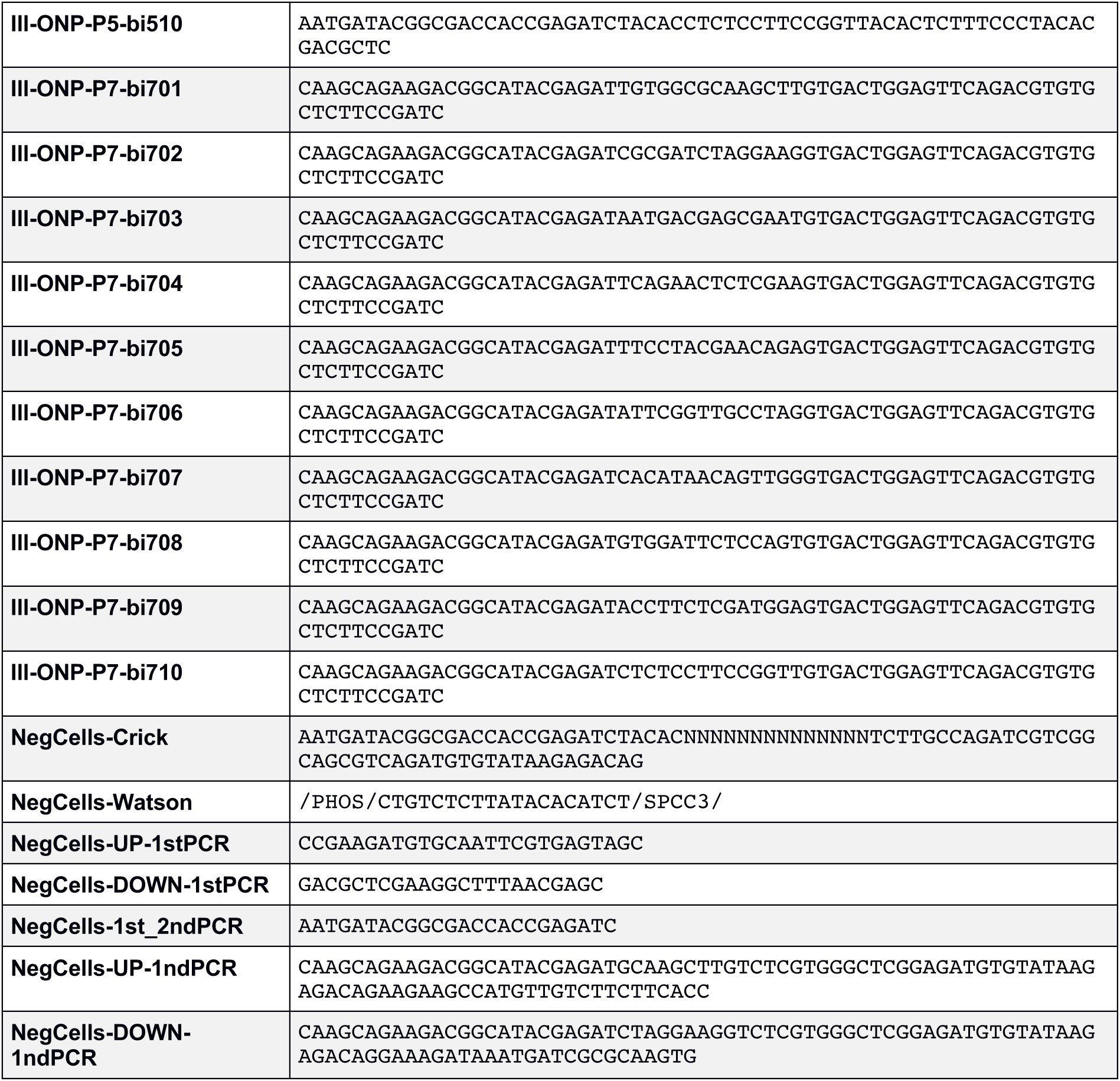
Primers used in this study.

**Supplementary Table 7.** Pool designs.

| Oligo-pool ID | PAM space | degeneracy |  | oligo-nucleotide length | supplier |
| --- | --- | --- | --- | --- | --- |
|  |  | sequences | ORFs |  |  |
| pool A | TTV (1,577) | 1,577 | 215 nuclear ORFs | 160 nt | CustomArray Inc. |
| pool B1 |  |  |  |  | Twist Bioscience |
| pool B2 |  |  |  |  | Twist Bioscience |
| pool C | TTV (11,155)<br>+TYN (1,317) | 12,472 | 5,664 | 170 nt | CustomArray Inc. |
| pool D | TTV (27,000) | 27,000 | 5,940 | 160 nt | Agilent Technologies |

### Supplementary Data

**Supplementary Data 1.** 1,577 oligonucleotide pool sequences for the small library targeting 215 nuclear proteins with nuclear localization.

Data provided as Online Supplementary Material.

**Supplementary Data 2.** Oligonucleotide pool sequences for the first genome-wide library by ORF. Some of the unique set of 12,472 sequences can target more than a single ORF, which is why a total number of 12,514 entries is provided. We excluded seven entries for *YEL020W-A*, *YEL020C-B*, *YEL021W*, and *YEL022W*, which are near to *ura3*-52, the locus at which the Cas12a-family proteins were integrated in this study (Supplementary Note 2).

Data provided as Online Supplementary Material.

**Supplementary Table 3.** Oligonucleotide pool sequences for the second genome-wide library by ORF. Some of the unique set of 27,000 sequences can target more than a single ORF, which is why a total number of 27,640 entries is provided. We excluded 15 entries for *YEL020W-A*, *YEL020C-B*, *YEL021W*, and *YEL022W*, which are near to *ura3*-52, the locus at which the Cas12a-family proteins were integrated in this study (Supplementary Note 2).

Data provided as Online Supplementary Material.

